# In-cell residue-resolved NMR of micromolar α-synuclein and tau at 310K

**DOI:** 10.1101/2024.10.07.617093

**Authors:** Hélène Chérot, Théophile Pred’homme, Francois-Xavier Theillet

## Abstract

Aggregates of non-globular proteins are associated to several degenerative disorders, e.g. α-synuclein and tau involved in Parkinson’s and Alzheimer’s diseases. Do these proteins suffer progressive changes in conformations and interactions in pathologic situations? In-cell NMR provides atomic-scale information in live cells but, until now, only at ~283 K in the case of unfolded proteins. Here, we report new labeling and acquisition methods enabling in-cell NMR at 310 K to study these proteins at micromolar concentrations, i.e. native cellular abundances. We used stable human cell lines expressing α-synuclein or tau upon induction in a culture medium supplemented with ^13^C-labeled amino acids, or precursors thereof. Acquiring ^13^Cα-^13^CO spectra permitted an early residue-resolved analysis of α-synuclein and tau at 310 K and <10 μM in HEK cells at 700 MHz. We detected disordered conformations and identical patterns of cellular interactions for α-synuclein wild-type and two mutants (F4A, A30P). Only the disordered N-terminus of tau was observable, even upon microtubule dismantling by colchicin. Our approach offers an excellent scalability -in signal and resolution-up to 1.2 GHz. ^13^C-labeling and ^13^C-detected NMR in live human cells are thus viable techniques for in-cell structural biology.

## Introduction

In-cell structural biology provides information on the conformational behaviors and binding abilities of proteins or nucleic acids in cellular milieus. This field is emerging thanks to the development of a range of complementary techniques, including cryo-electron tomography, EPR, mass-spectrometry, NMR, or FRET ^[1–9]^. In-cell NMR exploits isotope-filters to observe selectively ^13^C-,^15^N-or ^19^F-labeled peptides (or nucleic acids) either delivered or transiently expressed in cells, which contain 1% or less of these isotopes at natural abundance ^[1,10]^.

NMR spectroscopy has notably the unique capacity to extract atomic-scale information on local structures and interactions of intrinsically disordered (regions of) proteins (IDRs/IDPs) ^[11,12]^, a class of natively non-folded peptides representing about 30% of eukaryotic proteomes and a great variety of key functions in cells ^[13–15]^. IDPs lack stable 3D structures, which makes them malleable objects ^[15,16]^, whose structural behavior in cells is thus to be questioned ^[17,18]^. This is especially true for those IDPs, whose misfolded forms are important actors of neurodegenerative disorders, like α-synuclein (α-syn) and tau ^[19–21]^.

We and others reported atomic-scale studies on IDPs using in-cell NMR, showing that unfolded states can be stable in human cells ^[1,22–24]^. However, these analyses were carried out in live cells maintained at 283 K or less: they were based on the observation of backbone amide ^1^H-^15^N NMR signals of ^15^N-labeled IDPs, signals that weaken and overlap severely at physiological pH and temperature, owing to fast water-amide ^1^H-exchange (>25 Hz) ^[11,25–28]^. This is unfortunate, because IDPs’ conformational ensembles are temperature dependent ^[29–31]^, and molecular activities of human cells are obviously limited at 283 K.

Other NMR approaches have been developed recently, which are immune to water-amide ^1^H exchange, giving access to residue-specific information on IDPs in physiological conditions ^[32–35]^. Among these, 2D ^13^Cα-^13^CO correlation experiments appeared the most suited to in-cell NMR studies: i) they would not be affected by the broad cellular ^1^H_2_O signal, ii) they would permit a clean isotope-filter, and iii) they would offer decent signal-to-noise ratios for IDPs. The feasibility of using ^13^Cα-^13^CO in-cell NMR experiments was still questionable in the absence of commercial ^13^C-labeling culture media or NMR-adapted ^13^C-labeling protocols. To get closer to genuine cellular conditions, we set out to establish a consistent approach to generate and analyze in-cell NMR samples, using stable-inducible cell lines, homemade culture media supplemented with isotope-labeled amino acids, and ^13^Cα-^13^CO NMR. We present below the step tests of this in-cell NMR scheme, and demonstrate its feasibility and potential usefulness.

## Results and Discussion

### Stable HEK cell lines with inducible expression of α-synuclein or tau

We sought to set up an in-cell NMR approach using protein expression *in situ*, avoiding the nowadays more popular delivery of purified isotope-labeled material ^[1]^. We wanted to avoid transient transfection too, which provokes broadly inhomogeneous cell populations in regard to protein expression (Figure 1a). We aimed at expressing proteins of interest in an inducible fashion: this would allow temporary isotope-labeling during protein expression, which would minimize the NMR signal from the cellular background. We chose to use a commercial HEK cell line, namely Flp-In™ T-Rex™ 293, which permits to insert a gene of interest (GOI) in a single transcriptionally active locus of the genome under the control of a doxycycline-regulated, hybrid CMV/TetO_2_ promoter (Figure 1b). This enables a homogeneous strong expression (Figure 1a) upon supplementation with doxycycline at 10 ng/mL (Figure S1a), which is way below the ~5 μg/mL necessary to interfere with exogenous α-synuclein aggregation in cells ^[36]^. We selected stable cell pools using the adapted antibiotics, and obtained non-clonal inducible cell lines for various α-syn and tau constructs. These expressed very reproducible quantities after 48 h of exposure to doxycycline. The intracellular concentrations reached 9.5 μM for α-syn (Figure 1c) and 7.5 μM for tau (Figure S1c), i.e. in the range of native concentrations, i.e. ~40 and ~5 μM, respectively ^[37,38]^ (or ~3 and ~1 copies per 1000 protein molecules in the human brain according to https://pax-db.org/ ^[39]^). The molecular content of these cells was also reproducible, hence generating a constant cellular background NMR signal. This enabled us to obtain background-free spectra, by subtracting spectra recorded with non-induced cells to those recorded with induced cells (Figure 1d).

**Figure 1.**
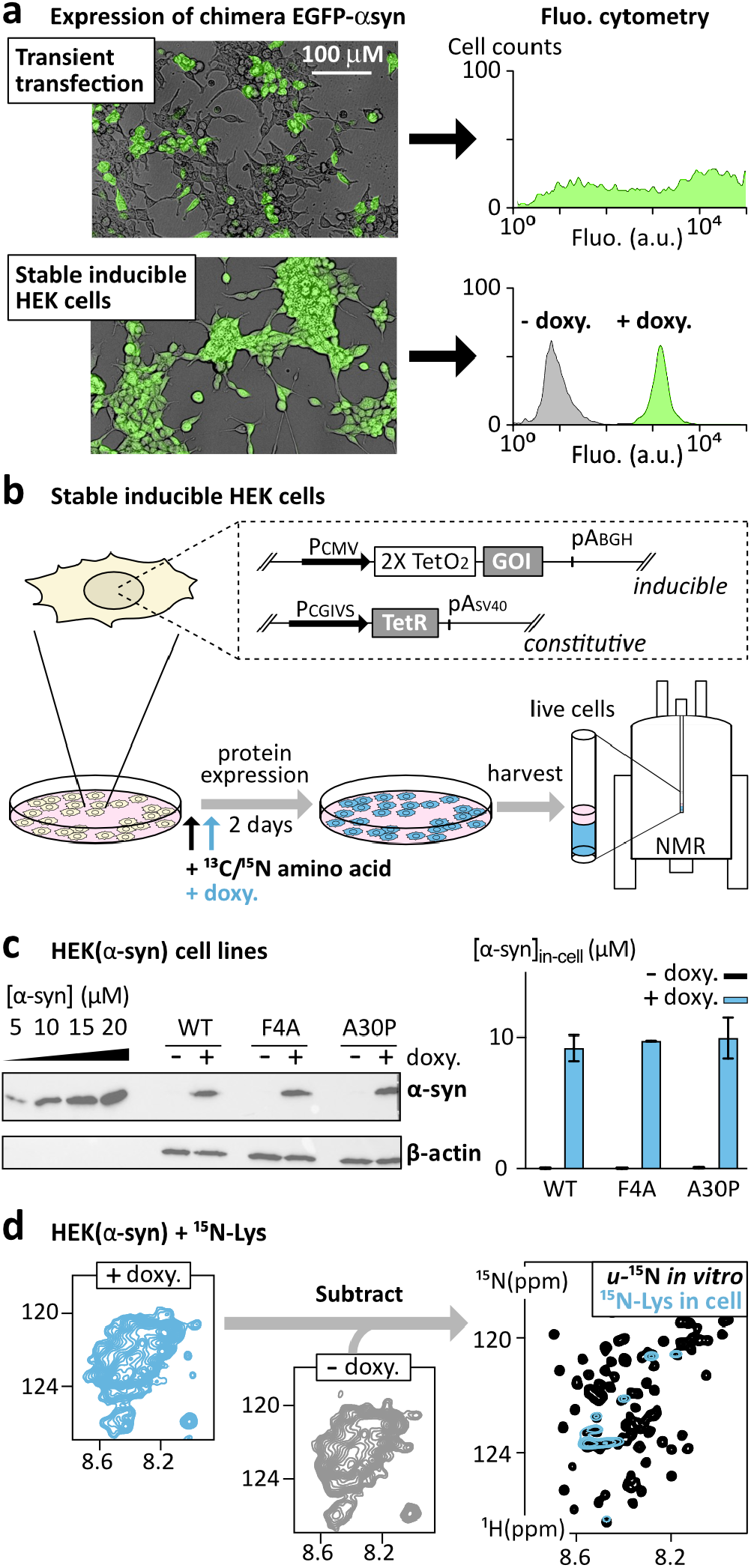
Principles of the sample preparation. **(a)** Expression of a chimera construct EGFP-αsyn in HEK cells using transient transfection or a stable inducible HEK cell line (single locus insert): *left*: overlay of brightfield and fluorescence microscopy; *right:* fluo-cytometry from transfected HEK cells (*up*) and non-induced (-doxycycline, grey) or induced (+doxy., green) stable inducible HEK cells (*down*). **(b)** Our stable-inducible HEK cell lines contain genes of interest (GOIs) inserted in a single chromosomal locus repressed by TetR on the *tet* operator 2 TetO2; TetR is released upon binding to doxy., which triggers GOI expression; cells grow in a standard medium, are transferred into a homemade medium containing selected ^13^C- and/or ^15^N-labeled amino acids, doxy.-induced 4 h later, and incubated during 48 h before harvesting. **(c)** Semi-quantitative western-blotting of α-syn showing comparable protein levels in cell lines expressing α-syn WT, F4A or A30P; the histogram shows results from triplicates. **(d)** Spectra recorded with non-induced cells serve to subtract the cellular background signals from natural abundance ^13^C/^15^N-species or from those produced during the incubation in presence of ^13^C/^15^N amino acids.

### Amino acid-specific isotope labeling of stable-inducible HEK cells

HEK cells are standardly cultured in a classical DMEM medium, whose recipe is public. Hence, we could emulate it and control the content in amino acids, eventually ^13^C- and/or ^15^N-labeled (Table S1). Earlier in-cell NMR studies have achieved protein isotope-labeling in mammalian cells using uniform ^15^N-labeling, which is conveniently obtained using commercial culture media (~100-200€ per sample) ^[1,10]^. Amino acid specific ^13^C-/^15^N-labeling could be a cheaper approach, also avoiding peak overlaps in IDPs, which we sought to test on our stable-inducible cell lines. A number of metabolic pathways are off in human cells (https://www.genome.jp/pathway/hsa01230), which might give access to novel labeling schemes for NMR.

We started with cells expressing α-syn, because it yields intense

NMR signals. We converged to the following experimental scheme: we switch to a culture medium containing selected ^13^C-/^15^N-amino acids 4 hours before inducing α-syn expression, and then incubate 48 hours before harvesting cells. We carried out successfully ^15^N-labeling of Asn, Lys, Phe, Tyr residues without any marked scrambling in 2D ^1^H-^15^N HSQC in-cell NMR spectra at 283 K (Figure 2a). We also succeeded in incorporating ^15^N-Gly in α-syn, although Gly scrambles partially with Ser amino acids. This pushed us to remove Ser from the culture medium, which resulted in a robust ^15^N-Gly and partial ^15^N-Ser labeling. After subtraction of a spectrum from non-induced cells, the observed crosspeaks have the chemical shifts of purified α-syn, with intensities modulated by the in-cell environment (see below). At the opposite, supplementing the culture medium with ^15^N-Asp, - Leu, -Val or -Ile yielded a wide spread of amide ^15^N signals, i.e. no selective labeling of those amino acids (Figure S2). These results are consistent with the common ^2^D/^13^C/^15^N-Lys/Arg labeling in SILAC-MS for quantitative proteomics ^[40]^, and with the handful of NMR-related reports about amino acid specific ^15^N-labeling in transiently transfected mammalian cells ^[41–44]^.

**Figure 2.**
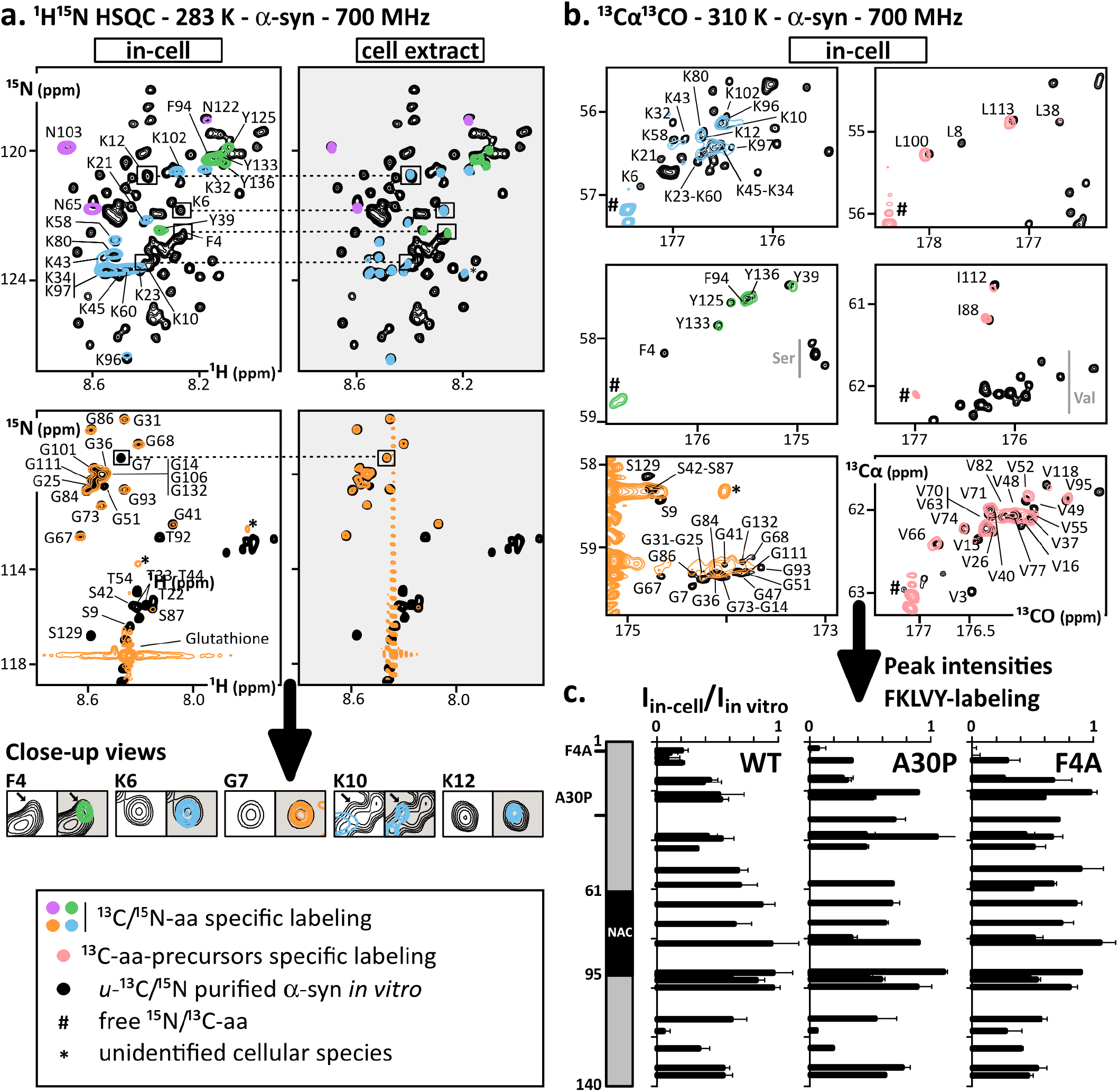
Amino acid specific isotope labelling and analysis of -syn in-cell. **(a)** Overlay of 2D ^1^H-^15^N HSQC spectra of purified [*u*-^15^N] -syn (black) and of cells expressing -syn in presence of ^15^N-Asn (purple), ^15^N-Lys (blue), ^15^N-Phe/^15^N-Tyr (green) or ^15^N-Gly (yellow) (spectra from non-induced cells were subtracted); spectra were recorded on intact cells or in clarified cell extracts (after sonication and a brief boiling step). **(b)** Overlay of 2D ^13^C -^13^CO spectra of purified recombinant [*u*-^13^C] -syn (black) and of cells expressing -syn in presence of ^13^C-Lys (blue), ^13^C-Phe/^13^C-Tyr (green) or ^13^C-precursors of Leu, Val or Ile (green) (spectra from non-induced cells were subtracted). **(c)** Residue-specific peak intensity ratios in ^13^C -^13^CO spectra of cells expressing ^13^C-F/K/L/V/Y labeled -syn WT/A30P/F4A versus purified -syn (I_*in-cell*_/I_*in vitroI*_) along the primary structure at 310 K.

Also consistent, we observed efficient incorporations of ^13^C-Asn, -Lys, -Phe, -Tyr with no scrambling issues, and of ^13^C-Glyresulting also in partial ^13^C-Ser labeling. These permitted the first ^13^C-detected in-cell protein NMR spectra in human cells at 310 K, using 4 hours-long ^13^Cα-^13^CO experiments at 700 MHz (Figure 2b, Figure S3). These revealed crosspeaks corresponding to those of purified, isolated α-syn *in vitro*. Interestingly, we did not have to carry out any background subtraction for ^13^C-Phe/Tyr, the cellular signal being below the noise level. Then, building on the observation that amide nitrogen atoms of Leu/Val/Ile were readily exchanged and diluted in the cellular content, we thought to use their ^13^C-labeled precursors α-ketoisocaproate, α-ketoisovalorate and 2-keto-3-methylvalerate, respectively (Figure S2). These cheaper alternatives were effectively converted to their amino acid counterpart, and permitted to record in-cell NMR ^13^Cα-^13^CO spectra of α-syn at 310 K, which showed exclusively Leu, Val or Ile signals (Figure 2b, Figure S3). These results are in agreement with very recent labeling studies on transiently transfected suspension HEK cells ^[45–47]^. 2D ^1^H-^13^C HSQC spectra revealed that these ^13^C-amino acids, or precursors thereof, were not or very weakly processed into undesired metabolites (Table S2), except for ^13^C-Gly incorporated in glutathione as already reported ^[48]^.

Altogether, we succeeded in incorporating a number of ^13^C-or ^15^N-amino acids during the expression of a GOI in stable-inducible HEK cells. This incorporation suffers almost no scrambling, which allows recording in-cell NMR spectra. It requires only mg quantities of the individual amino acids, which cost only ~1 to 10€ per sample depending on the amino acid (Table S1).

### Investigating α-syn at 310 K in cells using ^13^Cα^13^CO

First, we noticed in 2D ^1^H-^15^N HSQC recorded at 283 K that α-syn expressed *in situ* at ~10 μM produced NMR signals similar to those observed in published studies, where recombinant purified α-syn had been delivered in human cells using electroporation ^[22,23]^. The crosspeaks are those from a monomeric, disordered α-syn. We did not detect its N-terminal residues (residues 1-20) in live cells but crosspeaks of the N-ter acetylated form of α-syn reappeared in heated cell extracts (Figure 2a). Indeed, α-syn is heat stable ^[49]^ (Figure S4), and thus the peak disappearance in cells revealed interactions with heat-precipitable cellular components, among which chaperones Hsc70 and Hsp90 play a major role according to previous studies ^[22,23]^. However, α-syn’s interactions with chaperones and lipid membranes are temperature dependent ^[23,50,51]^. These can now be studied at physiological pH and temperature in a residue-resolved fashion using ^13^Cα-^13^CO experiments. As a test case, we recorded spectra of α-syn in presence of large unilamellar vesicles (LUVs) made from pig brain polar lipids: we observed lower signal intensities revealing an interaction with the LUVs on the first 100 amino acids only at 310 K, and not at 283 K (Figure S5); this was thus not detectable using the classical ^1^H-^15^N HSQC.

We sought to investigate this aspect at 310 K in ^13^C-K/F/L/V/Y-labeled cells expressing α-syn. First, the detected signals overlapped with those of the purified α-syn (Figure S6, Figure S7 shows the absence of leakage). However, most of the ^13^Cα-^13^CO crosspeaks of α-syn in cells were attenuated (Figure 2c). The profile of residue-resolved intensities looked like those observed in previous studies at 283 K: the 20 first residues were almost not detected, those neighboring L38-Y39 and the last 20 residues have attenuated intensities, all of which can be imputed to interactions with chaperones ^[22,23]^. We performed the same experiments with stable inducible cell lines expressing the mutants α-syn-F4A and the pathogenic α-syn-A30P, which both have reduced affinities to lipid membranes ^[22,52–54]^. We observed similar intensity profiles to those of α-syn-WT, which was consistent with an absence of binding to lipids (Figure 2c).

Altogether, the in-cell spectra revealed the presence of a population of disordered α-syn that is not in interaction with lipids but most probably with cellular chaperones.

### Investigating tau at 310 K in cells using ^13^Cα^13^CO

Then, we sought to apply our approach to other IDPs, and chose an emblematic long one, tau-2N4R (441 residues), whose role in so-called tauopathies calls for improved knowledge ^[21]^. Because earlier assignments were obtained on shorter constructs or at pH 6.5 or less, we achieved a near-complete backbone assignment at pH 7 and 283 K using a 1.2 GHz spectrometer (BMRB 52554). We transferred the ^13^Cα-^13^CO assignment at 310 K using a temperature gradient (Figure S8). Then, we recorded ^13^Cα-^13^CO spectra -at 310 K and 700 MHz-of a stable inducible cell line expressing tau-2N4R, which we labeled with ^13^C-F/I/L/R/V/Y. To obtain exploitable NMR signals, we had to supplement the culture medium with butyric acid at 2 mM, which increases the doxycycline-induced expression by a factor 2 (Figure S1b). We observed only a few weak crosspeaks overlapping with those from the ~150 N-terminal residues of purified tau (Figure 3a). At the opposite, other peaks were reproducibly missing, notably those from residues in the R1-R2-R3-R4 region (Figure 3b). This may reveal a population of tau that is free from any interaction in the N-terminal region, while the microtubule (MT) binding regions are either adopting ordered structures and/or transient interactions with cellular components. These regions are known to bind chaperones and MTs, and also to be involved in aggregates found in patients with tauopathies ^[21,55,56]^. To test the hypothesis of MTs binding, we supplemented tau-expressing HEK cells with colchicine at 20 μM during 24 h, leading to MTs disassembly and the progressive detachment of cells from the flask. In-cell spectra were highly similar to those of non-treated cells, showing only weak signals from the N-terminal residues, possibly until aa200. Hence, our results do not seem to reveal a dominant binding to MTs, but fit better to tau-2N4R binding to chaperones or being involved in liquid-liquid phase separation ^[57]^.

**Figure 3:**
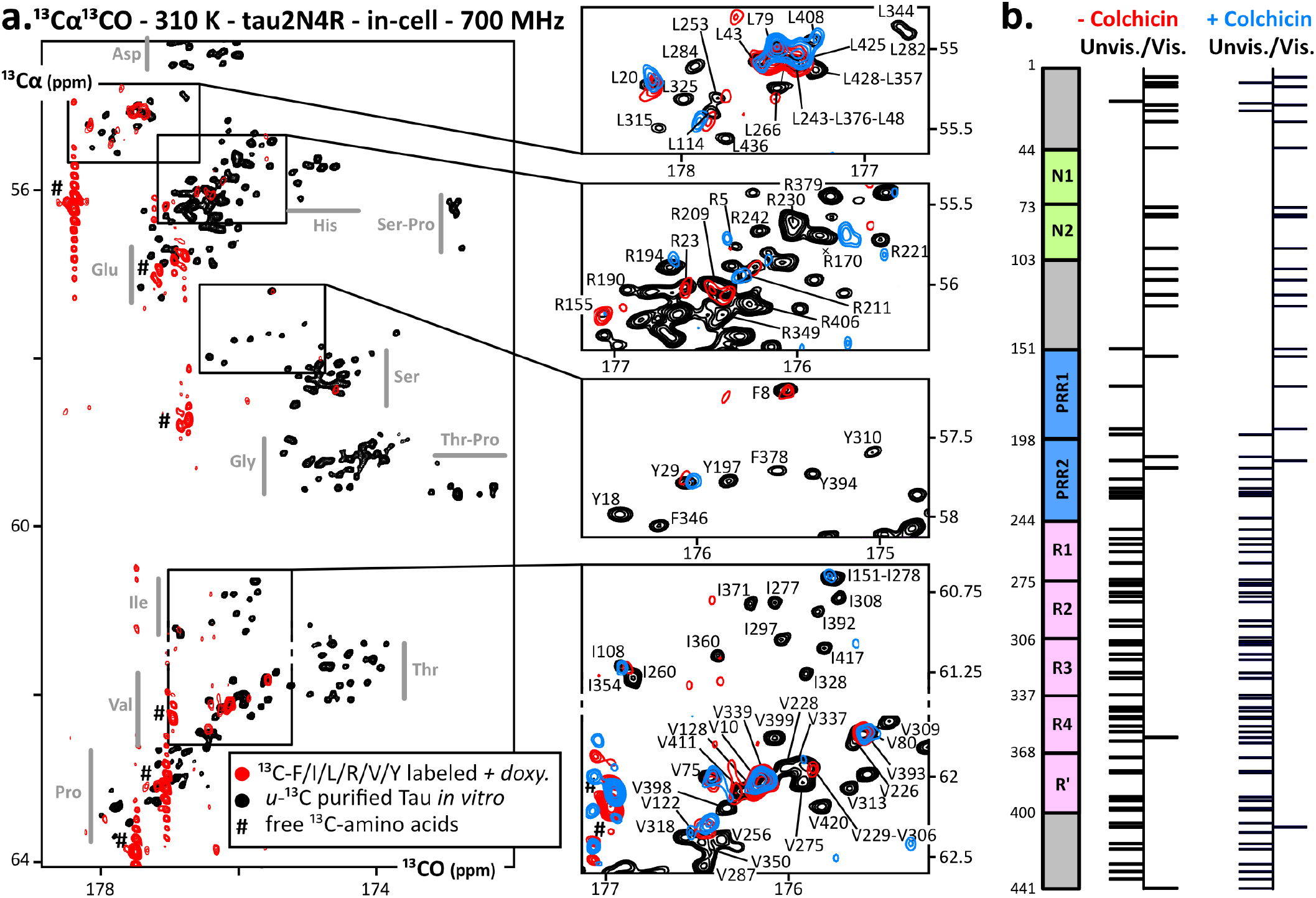
In-cell NMR of tau2N4R expressed in situ. **(a)** Overlay of 2D ^13^C -^13^CO spectra of purified recombinant [*u*-^13^C] tau (black) and of cells expressing ^13^C-F/I/L/R/V/Y tau in absence (red) or in presence of colchicine (blue) (spectra from non-induced cells were subtracted); cells use ^13^C-Arg to produce some ^13^C-Pro. **(b)** Scheme of tau2N4R primary structure (441 residues) and summary of peaks that are unambiguously detected or non-detected.

### ^13^Cα^13^CO acquisition at higher magnetic fields

We verified that ^13^Cα-^13^CO experiments can provide improved signal and resolution up to 1.2 GHz, the highest commercial magnetic field accessible. It was questionable because amide ^13^CO have a large chemical shift anisotropy (CSA), which provokes T2 relaxation scaling with the square of the magnetic field ^[58–60]^. This would come with broader and weaker signals at ultrahigh fields, counterbalancing the desired improvements of large magnets on resolution and signal. Interestingly, we measured almost constant ^13^CO linewidths, and peak intensities scaling with the probe sensitivity from 700 to 1200 MHz (Figure S9). This is in agreement with recent evaluations of ^13^C-detected experiments on IDPs at 1.2 GHz ^[61]^. It is consistent with the facts that i) IDPs are weakly affected by CSA effects due to their high flexibility and ii) ^13^C-detection sensitivity scales well with the magnetic field even in watery and salty samples analyzed using a cryoprobe ^[62]^. We had an immediate access to a 700 MHz spectrometer equipped with a ^13^C-detection optimized probe, which offered a much higher ^13^C-sensitivity than the accessible 950 and 1200 MHz spectrometers equipped with ^1^H-optimized probes. Using a ^13^C-dedicated probe at 1.2 GHz should provide about twice the resolution and sensitivity of our 700 MHz spectrometer for ^13^Cα^13^CO experiments.

## Conclusion

Our work aimed at establishing experimental conditions for in-cell NMR of IDPs closer to physiological conditions. The present set of methods proved to give access to residue specific information on α-syn and tau at intracellular concentrations of ~10 μM and at 310 K. Even though intrinsically less sensitive than ^1^H-detection, ^13^C-detection permits to approach such native concentrations. It has great advantages for in-cell analysis: ^13^C-detection is much less affected by water signal and inhomogeneities in magnetic susceptibility than ^1^H-detection ^[63,64]^.

From the technical point of view, a few points can be highlighted. Leucine, valine and isoleucine could be incorporated using their immediate ketoacids precursors, which are subject to a high transaminase activity, in agreement with recent reports ^[45,47]^. This will permit ^13^C-labeling at a reduced price in mammalian cells. At the opposite, some amino acids could be incorporated without any ^15^N-scrambling, in agreement with other studies ^[41–44]^. This should enable residue specific ^13^C/^15^N-combined to ^2^H-labeling schemes in mammalian cells, offering high sensitivity on folded proteins ^[47,65]^.

From the biological point of view, we detected populations of α-syn and tau adopting disordered conformations in cells at 310 K. α-syn appears to interact with cellular species on its N- and C-termini, in a similar fashion than at 283 K. Concerning tau, the treatment by colchicine revealed that the missing signals between residues 150 and 441 of tau were not or not only due to microtubule binding. These early analyses motivate further investigations, and will most probably require the complementary use of solution and solid-state NMR.

Future studies shall examine improved cellular models: HEK cells are convenient to manipulate, but neuronal cells would be more relevant to study α-syn/tau-linked neurodegeneration. The use of a flow-probe bioreactor will also be instrumental to achieve a longitudinal monitoring in steady, wealthy conditions ^[10]^. This requires to trap cells in gels, which decreases the number of detected molecules in the NMR probe, hence their NMR signal. Using our ^13^Cα^13^CO approach, an intracellular concentration of 10 μM is a lower limit to obtain exploitable signals in a few hours at 700 MHz. Higher fields will help solving this sensitivity issue: the ^13^Cα^13^CO experiment yields S/N scaling with the field, in contrast to ^1^H-detected experiments ^[62]^. Higher cellular concentrations of α-syn/tau, i.e. ~20 μM, would yield much better spectra, while remaining in the range of native concentrations for α-syn and tau. This can be reached by multiple insertions of the GOI ^[66–68]^.

Even though recent methods have been developed to predict isolated IDPs conformational ensembles ^[16,69]^, experimental information on the effects of cellular conditions are still needed to understand how IDPs behave, function and eventually misfold. We presented here a set of methods that will enable such experimental studies, providing residue specific in-cell characterizations of IDPs at physiological concentrations and temperatures.

## Supporting information

SuppInfo

## Supporting Information

The authors have cited additional references within the Supporting Information.

## Acknowledgements

This work was supported by the CNRS and the CEA-Saclay (CEA/PSAC/DPRS/BE/TG/2021-410), by the French Infrastructure for Integrated Structural Biology (https://frisbi.eu/, grant number ANR-10-INSB-05-01, Acronym FRISBI) and by the French National Research Agency (ANR; research grants ANR-14-ACHN-0015 and ANR-20-CE92-0013). Financial support from the IR INFRANALYTICS FR2054 for conducting the research is gratefully acknowledged, and we value the commitment and expertise of X. Trivelli and F.-X. Cantrelle.

