## Supplementary material for "In-cell residue-resolved NMR of micromolar α-synuclein and tau at 310K": SuppInfo

Supporting information for this article is given via a link at the end of the document.

**Abstract:** Aggregates of non-globular proteins are associated to several degenerative disorders, e.g. $\alpha$-synuclein and tau involved in Parkinson's and Alzheimer's diseases. Do these proteins suffer progressive changes in conformations and interactions in pathologic situations? In-cell NMR provides atomic-scale information in live cells but, until now, only at ~283 K in the case of unfolded proteins. Here, we report new labeling and acquisition methods enabling in-cell NMR at 310 K to study these proteins at micromolar concentrations, i.e. native cellular abundances. We used stable human cell lines expressing $\alpha$-synuclein or tau upon induction in a culture medium supplemented with $^{13}$C-labeled amino acids, or precursors thereof. Acquiring $^{13}$C$\alpha$-$^{13}$CO spectra permitted an early residue-resolved analysis of $\alpha$-synuclein and tau at 310 K and <10 $\mu$M in HEK cells at 700 MHz. We detected disordered conformations and identical patterns of cellular interactions for $\alpha$-synuclein wild-type and two mutants (F4A, A30P). Only the disordered N-terminus of tau was observable, even upon microtubule dismantling by colchicin. Our approach offers an excellent scalability -in signal and resolution- up to 1.2 GHz. $^{13}$C-labeling and $^{13}$C-detected NMR in live human cells are thus viable techniques for in-cell structural biology.

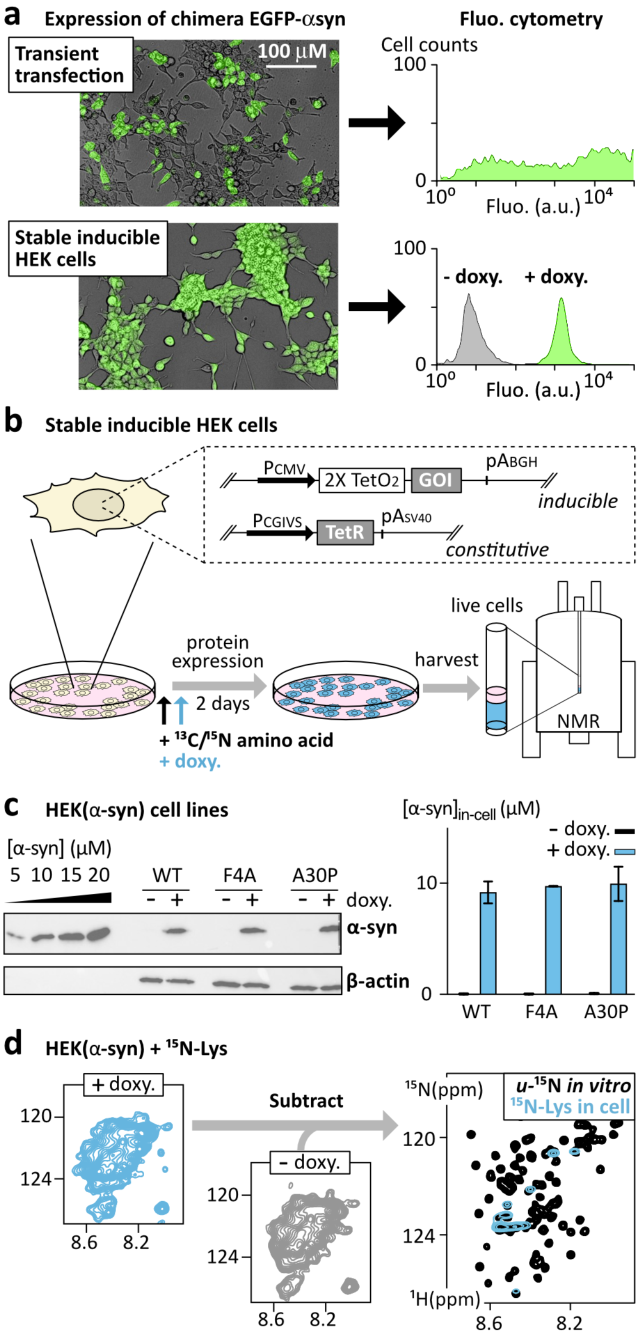


**Figure 1. Principles of the sample preparation. (a)** Expression of a chimera construct EGFP-αsyn in HEK cells using transient transfection or a stable inducible HEK cell line (single locus insert): *left*: overlay of brightfield and fluorescence microscopy; *right:* fluo-cytometry from transfected HEK cells (*up*) and non-induced (-doxycycline, grey) or induced (+doxy., green) stable inducible HEK cells (*down*). **(b)** Our stable-inducible HEK cell lines contain genes of interest (GOIs) inserted in a single chromosomal locus repressed by TetR on the *tet* operator 2 $TetO_2$; TetR is released upon binding to doxy., which triggers GOI expression; cells grow in a standard medium, are transferred into a homemade medium containing selected $^{13}C$- and/or $^{15}N$-labeled amino acids, doxy.-induced 4 h later, and incubated during 48 h before harvesting. **(c)** Semi-quantitative western-blotting of α-syn showing comparable protein levels in cell lines expressing α-syn WT, F4A or A30P; the histogram shows results from triplicates. **(d)** Spectra recorded with non-induced cells serve to subtract the cellular background signals from natural abundance $^{13}C/^{15}N$-species or from those produced during the incubation in presence of $^{13}C/^{15}N$ amino acids.

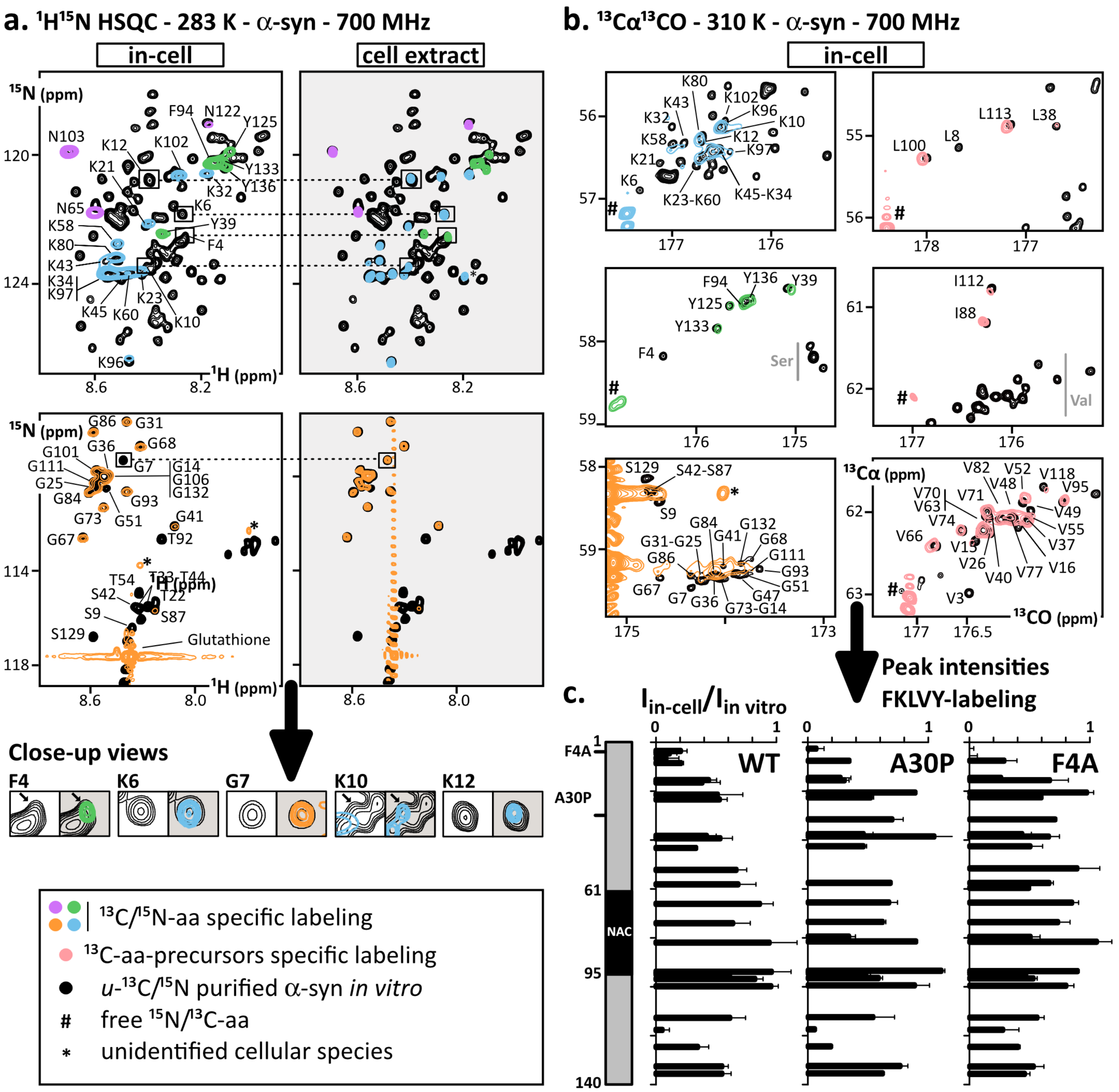


**Figure 2. Amino acid specific isotope labelling and analysis of α-syn in-cell. (a)** Overlay of 2D $^{1}$H-$^{15}$N HSQC spectra of purified [*u*-$^{15}$N] α-syn (black) and of cells expressing α-syn in presence of $^{15}$N-Asn (purple), $^{15}$N-Lys (blue), $^{15}$N-Phe/$^{15}$N-Tyr (green) or $^{15}$N-Gly (yellow) (spectra from non-induced cells were subtracted); spectra were recorded on intact cells or in clarified cell extracts (after sonication and a brief boiling step). **(b)** Overlay of 2D $^{13}$Cα-$^{13}$CO spectra of purified recombinant [*u*-$^{13}$C] α-syn (black) and of cells expressing α-syn in presence of $^{13}$C-Lys (blue), $^{13}$C-Phe/$^{13}$C-Tyr (green) or $^{13}$C-precursors of Leu, Val or Ile (green) (spectra from non-induced cells were subtracted). **(c)** Residue-specific peak intensity ratios in $^{13}$Cα-$^{13}$CO spectra of cells expressing $^{13}$C-F/K/L/V/Y labeled α-syn WT/A30P/F4A versus purified α-syn ($I_{in\text{-}cell}/I_{in\ vitro}$) along the primary structure at 310 K.

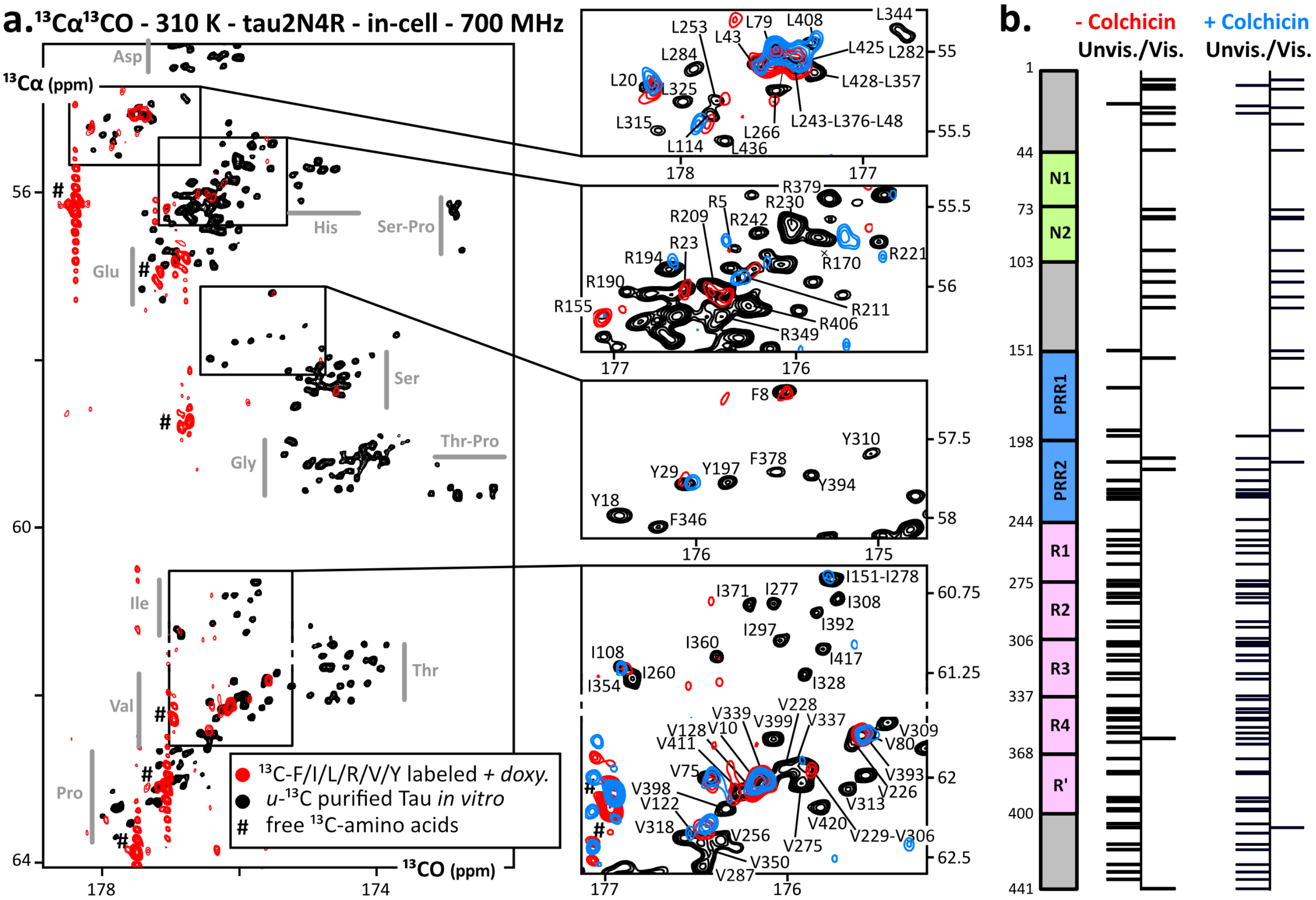


**Figure 4: In-cell NMR of tau2N4R expressed in situ. (a)** Overlay of 2D $^{13}$Cα-$^{13}$CO spectra of purified recombinant [*u*-$^{13}$C] tau (black) and of cells expressing $^{13}$C-F/I/L/R/V/Y tau in absence (red) or in presence of colchicine (blue) (spectra from non-induced cells were subtracted); cells use $^{13}$C-Arg to produce some $^{13}$C-Pro. **(b)** Scheme of tau2N4R primary structure (441 residues) and summary of peaks that are unambiguously detected or non-detected.

**Keywords:** In-cell NMR • Intrinsically Disordered Proteins • α-Synuclein • tau • Isotope labeling

[1] F.-X. Theillet, *Chem. Rev.* **2022**, *122*, 9497–9570.
[2] E. Nogales, J. Mahamid, *Nature* **2024**, *628*, 47–56.
[3] D. Goldfarb, *Curr. Opin. Struct. Biol.* **2022**, *75*, 102398.
[4] M. Yu, M. Heidari, S. Mikhaleva, P. S. Tan, S. Mingu, H. Ruan, C. D. Reinkemeier, A. Obarska-Kosinska, M. Siggel, M. Beck, G. Hummer, E. A. Lemke, *Nature* **2023**, *617*, 162–169.
[5] B. Schuler, I. König, A. Soranno, D. Nettels, *Angew. Chem. Int. Ed.* **2021**, anie.202016804.
[6] J. M. Plitzko, B. Schuler, P. Selenko, *Curr. Opin. Struct. Biol.* **2017**, *46*, 110–121.
[7] Z. Zhang, Q. Zhao, Z. Gong, R. Du, M. Liu, Y. Zhang, L. Zhang, C. Li, *JACS Au* **2024**, *4*, 369–383.
[8] C. L. McCafferty, S. Klumpe, R. E. Amaro, W. Kukulski, L. Collinson, B. D. Engel, *Cell* **2024**, *187*, 563–584.
[9] M. Beck, R. Covino, I. Hänelt, M. Müller-McNicoll, *Cell* **2024**, *187*, 545–562.
[10] E. Luchinat, M. Cremonini, L. Banci, *Chem. Rev.* **2022**, *122*, 9267–9306.
[11] H. J. Dyson, P. E. Wright, *Curr. Opin. Struct. Biol.* **2021**, *70*, 44–52.
[12] A. R. Camacho-Zarco, V. Schnapka, S. Guseva, A. Abyzov, W. Adamski, S. Milles, M. R. Jensen, L. Zidek, N. Salvi, M. Blackledge, *Chem. Rev.* **2022**, *122*, 9331–9356.
[13] E. A. Lemke, M. M. Babu, R. W. Kriwacki, T. Mittag, R. V. Pappu, P. E. Wright, J. D. Forman-Kay, *Mol. Cell* **2024**, *84*, 1188–1190.
[14] M. C. Aspromonte, M. V. Nugnes, F. Quaglia, A. Bouharoua, DisProt Consortium, V. Sagris, V. J. Promponas, A. Chasapi, E. Fichó, G. E. Balatti, G. Parisi, M. G. Buitrón, G. Erdos, M. Pajkos, Z. Dosztányi, L. Dobson, A. D. Conte, D. Clementel, E. Salladini, E. Leonardi, F. Kordevani, H. Ghafouri, L. G. T. Ku, A. M. Monzon, C. Ferrari, Z. Kálmán, J. F. Nilsson, J. Santos, C. Pintado-Grima, S. Ventura, V. Ács, R. Pancsa, M. G. Kulik, M. A. Andrade-Navarro, P. J. B. Pereira, S. Longhi, P. L. Mercier, J. Bergier, P. Tompa, T. Lazar, S. C. E. Tosatto, D. Piovesan, *Nucleic Acids Res.* **2024**, *52*, D434–D441.
[15] A. S. Holehouse, B. B. Kragelund, *Nat. Rev. Mol. Cell Biol.* **2024**, *25*, 187–211.
[16] G. Tesei, A. I. Trolle, N. Jonsson, J. Betz, F. E. Knudsen, F. Pesce, K. E. Johansson, K. Lindorff-Larsen, *Nature* **2024**, *626*, 897–904.
[17] F.-X. Theillet, A. Binolfi, T. Frembgen-Kesner, K. Hingorani, M. Sarkar, C. Kyne, C. Li, P. B. Crowley, L. Gierasch, G. J. Pielak, A. H. Elcock, A. Gershenson, P. Selenko, *Chem. Rev.* **2014**, *114*, 6661–6714.
[18] D. Moses, G. M. Ginell, A. S. Holehouse, S. Sukenik, *Trends Biochem. Sci.* **2023**, *48*, 1019–1034.
[19] L. M. A. Oliveira, T. Gasser, R. Edwards, M. Zweckstetter, R. Melki, L. Stefanis, H. A. Lashuel, D. Sulzer, K. Vekrellis, G. M. Halliday, J. J. Tomlinson, M. Schlossmacher, P. H. Jensen, J. Schulze-Hentrich, O. Riess, W. D. Hirst, O. El-Agnaf, B. Mollenhauer, P. Lansbury, T. F. Outeiro, *Npj Park Dis* **2021**, *7*, 65.
[20] T. Simuni, L. M. Chahine, K. Poston, M. Brumm, T. Buracchio, M. Campbell, S. Chowdhury, C. Coffey, L. Concha-Marambio, T. Dam, P. DiBiaso, T. Foroud, M. Frasier, C. Gochanour, D. Jennings, K. Kieburtz, C. M. Kopil, K. Merchant, B. Mollenhauer, T. Montine, K. Nudelman, G. Pagano, J. Seibyl, T. Sherer, A. Singleton, D. Stephenson, M. Stern, C. Soto, C. M. Tanner, E. Tolosa, D. Weintraub, Y. Xiao, A. Siderowf, B. Dunn, K. Marek, *Lancet Neurol.* **2024**, *23*, 178–190.
[21] B. C. Creekmore, R. Watanabe, E. B. Lee, *Annu. Rev. Pathol. Mech. Dis.* **2024**, *19*, 345–370.
[22] F.-X. Theillet, A. Binolfi, B. Bekei, A. Martorana, H. M. Rose, M. Stuiver, S. Verzini, D. Lorenz, M. van Rossum, D. Goldfarb, P. Selenko, *Nature* **2016**, *530*, 45–50.
[23] B. M. Burmann, J. A. Gerez, I. Matečko-Burmann, S. Campioni, P. Kumari, D. Ghosh, A. Mazur, E. E. Aspholm, D. Šulskis, M. Wawrzyniuk, T. Bock, A. Schmidt, S. G. D. Rüdiger, R. Riek, S. Hiller, *Nature* **2019**, *577*, 127–132.
[24] S. Zhang, C. Wang, J. Lu, X. Ma, Z. Liu, D. Li, Z. Liu, C. Liu, *Int J Mol Sci* **2019**, *20*, 90–14.
[25] S.-T. D. Hsu, C. W. Bertoncini, C. M. Dobson, *J. Am. Chem. Soc.* **2009**, *131*, 7222–7223.
[26] S. Kim, K.-P. Wu, J. Baum, *J. Biomol. NMR* **2013**, *55*, 249–256.
[27] T. Yuwen, N. R. Skrynnikov, *J Biomol NMR* **2014**, *58*, 175–192.
[28] Y. Bai, J. S. Milne, L. Mayne, S. W. Englander, *Proteins* **1993**, *17*, 75–86.
[29] R. Wuttke, H. Hofmann, D. Nettels, M. B. Borgia, J. Mittal, R. B. Best, B. Schuler, *Proc. Natl. Acad. Sci. U.S.A.* **2014**, *111*, 5213–5218.

### Supporting information

### In-cell residue-resolved NMR of micromolar $\alpha$-synuclein and tau at 310K

Hélène Chérot,[a] Théophile Pred'homme,[a] Francois-Xavier Theillet*[a,b]

[*] Corresponding author.

[a] H. Chérot, T. Pred'homme, Dr. F.X. Theillet, Université Paris-Saclay, CEA, CNRS, Institute for Integrative Biology of the Cell (I2BC), 91198, Gif-sur-Yvette, France.

[b] Université Paris Cité, CNRS, CiTCoM, F-75006 Paris, France


#### Table of Contents


| | |
|---|---|
| Experimental procedures | pages 2-5 |
| References | page 5 |
| Pulse sequence | pages 6-8 |
| Supplementary tables | pages 9-11 |
| Supplementary Figures | pages 11-20 |
| Authors contributions | page |


##### Recombinant production of [*u*-$^{13}C/^{15}N$]-peptides and their purification

α-syn and tau constructs were recombinantly expressed in *E. coli* BL21(DE3) cells (New England Biolabs, ref. C2527H), using pET-22b(+) (cloning sites NdeI-NotI) and pET-45b(+) (cloning sites KpnI-AvrII) plasmids, respectively. These contained cDNAs codon-optimized for bacterial expression synthesized and cloned by Genscript. Mutations were also executed by Genscript. α-Syn constructs were all N-terminally acetylated by yeast N-acetyltransferase complex B (NatB), which was co-expressed following the previously published protocols [1]. The tau constructs included a Tev cleavage site ENLYFQG separating the hexahistidine tag and the peptide fragments of interest. All cultures were induced at OD600 0.8 with isopropyl β-D-thiogalactopyranoside (IPTG, Euromedex) at 0.25 mM, for 4 h at 37 °C and overnight at 20 °C for α-Syn and tau constructs, respectively.
Non-isotope labeled proteins were produced in LB medium (Becton Dickinson ref. 244520). $^{15}N/^{13}C$ isotope-labelled proteins were obtained by overexpression in M9 minimal media supplemented with $^{15}NH_4Cl$ (0.5 g/L, Cortecnet) and *u*-$^{13}C$-glucose (2 g/L, Eurisotop-CIL).
Bacteria were pelleted by centrifugation (8 min at 4,000 g, room temperature). Monomeric α-syn was purified as described [1]. For tau constructs, the bacterial pellets were resuspended in 40 mL of a solution at pH=7.0 containing Tris at 20 mM, NaCl at 150 mM, DTT at 5 mM, PMSF at 1 mM, 500 μg of lysozyme, 0.5 μL of benzonase (Merck ref. E1014). Cells were sonicated on ice during 2.5 min using 1 s ON/ 4 s OFF cycles of sonication (50% amplitude). The lysates were clarified by centrifugation (15 minutes at 15,000 g at 4 °C). The soluble fraction was loaded on a Ni-NTA column (5 mL HisTrap FF, Cytiva) and the protein was eluted in TBS (Tris at 20 mM, NaCl at 150 mM, pH=7.5) using an imidazole gradient. The hexahistidine affinity tag was removed by Tev cleavage carried out 1 h at room temperature. Nucleic acids contaminants were separated from the protein using an anion exchange column (Resource Q, Cytiva). The elution fractions were concentrated to 2 mL and loaded on a gel filtration column (HiLoad 16/600 Superdex 200pg, Cytiva) equilibrated in PBS (phosphate at 20 mM, NaCl at 150 mM) at pH=7.0.

##### Generation of inducible stable cell lines

α-Syn and tau cDNAs were codon-optimized for expression in human cells and synthesized by Genscript, before being cloned into a pcDNA5/FRT/TO/EGFP vector (kind gift from A.M. Tassin) at BamHI/NotI restriction sites. The coding sequence for EGFP was removed from the parental plasmid by mutation (Genscript), and the later plasmid containing the genes coding for α-Syn-F4A, α-Syn-A30P and α-Syn-aa1-103 were also obtained by single point mutation (Genscript). The pOG44 Flp-Recombinase Expression Vector was purchased from Invitrogen (ref. V600520).
Parental Flp-In T-Rex HEK 293 cells were a kind gift from A.M. Tassin (Institute for Integrative Biology of the Cell, Gif-sur-Yvette, France). These were cultivated in Dulbecco's modified Eagle's medium (StableCell™ DMEM-high glucose-Glutamax, Sigma, ref. D0819) supplemented with 10% (v/v) fetal bovine serum (FBS, Gibco, ref. 10270-106) (abbrev. below: DMEM-FBS), 100 U/mL penicillin, 10 μg/mL streptomycin (Sigma, ref. P4333), 10 μg/mL blasticidin (Invivogen, Cat# ant-bl-1) and 100 μg/mL zeocin (Zeocin™ Selection Reagent, Gibco, ref. R25001).
For generating the stable cell lines, the parental Flp-In T-Rex HEK 293 cells were seeded in a 6-well plate (3.5 cm diameter), at a density of 300,000 cells per well and 24 h prior transfection. Cells were then washed with phosphate buffered saline (PBS, Sigma, D8537) and incubated 8 hours with a 1:1:2 ratio (w/w) of pcDNA5-(α-Syn/tau):pOG44:PEI in 2 mL of DMEM (using 4 μg of plasmid). The transfection reagent was PEI MAX™ (Transfection grade linear polyethyleiminine, MW 40,000, Polysciences ref. 24765-1). Cells were later washed and maintained in DMEM-FBS during 24 hours. Cells that integrated the α-Syn or tau cDNA were selected in DMEM-FBS supplemented with 200 μg/mL hygromycin, 100 U/mL penicillin, 10 μg/mL, streptomycin (Sigma, ref. P4333), 10 μg/mL blasticidin (Invivogen, Cat# ant-bl-1) and 100 μg/mL zeocin (Zeocin™ Selection Reagent, Gibco, ref. R25001). The medium was refreshed every day in the first 2 days, and every 5 days once the non-transformed cells detached under hygromycin pressure. Colonies were detected after 3 to 4 weeks. These were trypsinized and progressively transferred into larger culture plates to reach about 150 million cells (i.e. passage~20). These were finally trypsinized, washed and resuspended in FBS supplemented with 10% (v/v) DMSO (Sigma-Aldrich, ref. D8418). Aliquots of 5-10 million cells were slowly frozen at -70°C using a freezing container (Corning CoolCell LX), and stored in liquid nitrogen. They were later thawed and cultured in DMEM-FBS supplemented with 100 μg/mL hygromycin, 100 U/mL penicillin,10 μg/mL streptomycin (Sigma, ref. P4333), 5 μg/mL blasticidin (Invivogen, Cat# ant-bl-1) and 50 μg/mL zeocin (Zeocin™ Selection Reagent, Gibco, ref. R25001).

All cells were grown at 37°C, in humidified atmosphere at 5% $CO_2$. Cell lines were tested for mycoplasma contaminations and were found mycoplasma-free.

**In-cell NMR sample**

To generate in-cell NMR samples, cells from an established Flp-In T-Rex HEK 293 cell line were grown at 70% confluence in a 300 $cm^2$ tissue culture flask. Four hours before induction, cells were washed with PBS and the medium was replaced with homemade DMEM (according to the composition of DMEM-high glucose-Glutamax, Sigma, ref. D0819), supplemented with dialyzed FBS (Gibco, ref. A3382001), Glutamax (Gibco, ref. 35050-038) and with the desired labeled amino-acids (see Table S1). Cells were induced with doxycycline at 10 ng/mL, which we determined to be sufficient for maximal expression rates (Figure S1) (Sigma D9891). After 44 to 48 h of protein expression in these conditions, cells were detached using trypsin-EDTA (Sigma, ref. T4174), washed once with DMEM-FBS and then twice in PBS, resuspended in 450 μL of fresh DMEM-FBS supplemented with $D_2O$ at 10% v/v, and pelleted into a 5 mm (diameter) advanced Shigemi NMR.
For samples exposed to lipids, the cell culture medium was supplemented with oleic acid at 250 μM (Sigma ref. O1383) and cholesterol at 200 μM (Sigma ref. C3045) 24 hours after doxycycline treatment and 24 hours before harvesting. OA and CL were mixed in PBS at 2.5 and 2 mM, respectively, and immediately diluted in the culture medium.

**Verification of the absence of protein leakage**

Protein leakage was controlled after in-cell NMR acquisition by recording $^{13}C\alpha^{13}CO$ spectra of the sample supernatant, which was obtained as follows: cells were resuspended in the 450 μL of medium in excess in the NMR tube, transferred in a 1.5 mL tube, and pelleted by centrifugation at 100 g during 3 minutes; the resulting supernatant was analyzed using the same tube and NMR parameters than in-cell samples.

**Production of cell extracts for NMR analysis**

Lysis of HEK cells after in-cell NMR was performed by sonication on ice using Q700 sonicator (Qsonica). A wet pellet of cells (~300 μL) was sonicated on ice during 3 minutes using 5 s ON/ 25 s OFF cycles of sonication (40% amplitude) in 400 μL of PBS (Sigma ref. D1408), supplemented with protease inhibitors (cOmplete EDTA-free, Sigma ref. 05056489001) and DTT at 2 mM. The cell extracts were cleared using centrifugation at 15,000 g for 10 minutes at 4 °C. Folded proteins were precipitated by thermal denaturation at 95 °C for 3 minutes. The final cell extracts were obtained using centrifugation at 15,000 g for 10 minutes at 4 °C. NMR spectra were acquired on the supernatant.

**Western Blot and quantification of intracellular concentrations**

The protein expression quantification was performed on HEK cells after 48h in labeled DMEM with and without induction of protein expression. The cells were diluted to 25 million cells/mL and lysed in a RIPA lysis buffer (20 mM Tris, 150 mM NaCl, 1% (v/v) Nonidet® P40 (USB), 1% (v/v) sodium deoxycholate (Sigma-aldrich ref. D6750), 0,1% (v/v) SDS), supplemented with protease inhibitor (cOmplete, EDTA-free, Sigma ref. 05056489001) for 20 minutes on ice, followed by sonication on ice (5s, 40% amplitude using Q700 sonicator® Qsonica). Aliquots containing about 150 000 lysed cells were loaded per well for SDS-PAGE analysis (15% acrylamide). To quantify the concentration of α-Syn in these cells, a dilution series of 11-44 ng of purified α-Syn (expressed recombinantly in bacteria, see above) was also loaded on the gels. Assuming a cellular volume of 1 pL (average diameter 13±2 μm, Biorad TC20™ Automated Cell Counter), the corresponding intracellular concentrations of the latter series were 5, 10, 15 and 20 μM.
After SDS-PAGE migration, proteins were transferred onto nitrocellulose membranes (GE Healthcare Life Science), using Trans-Blot® turbo™ (Biorad). After blocking for 1h in 5% (w/v) skim milk (Sigma, ref. 70166) in TBST (0.1% (v/v) Tween-20, 20 mM Tris, 150 mM NaCl, pH 7.5), membranes were probed with the anti-α-Syn antibody ab-138501 (Abcam, 1:10,000 dilution), anti-tau HT-7 and T46 (Invitrogen, ref. MN1000 & 13-6400,1:1500 and 1:1000 dilutions, respectively) and anti-β-actine antibody A1978 (Sigma, 1:10,000 dilution) for 30 minutes to 1h at room temperature. Secondary antibodies were HRP-conjugated anti-rabbit or anti-mouse (Thermo Scientific, ref. 31460 and 31430, dilution 1:10,000), and were incubated during 45 minutes with the membranes. Membranes were developed using clarity Western ECL Substrate (BioRad). Luminescence signals were detected on a BioRad ChemiDoc™ imaging system, and quantified with Image Lab 5.1 (BioRad).

**NMR acquisition and processing of 2D reference and in-cell spectra**

In-cell NMR spectra were recorded using 700 MHz Bruker Avance Neo spectrometers, equipped with cryogenically cooled triple resonance probes, namely $^{1}H\{^{13}C/^{15}N\}$ TCI and $^{13}C$ $\{^{1}H/^{15}N\}$ TXO probes for spectra at 10°C ($^{1}H$-detection) and 37°C ($^{13}C$-detection), respectively. We used 5 mm Shigemi (for $^{1}H$-detection) or 5 mm Shigemi advanced (for $^{13}C$-detection) NMR tubes without plunger. All spectra were acquired and processed with Topspin 4. The TCI probe is from 2006, and its sensitivity was determined to be S/N=7260 for $^{1}H$ 0.1% ethylbenzene when delivered. The 700MHz-TXO probe is from 2020, its sensitivity was determined to be S/N=3400 for $^{13}C$ ASTM when delivered.
"cell-extract" and "in-cell" spectra were recorded with exactly the same parameters for the "induced" and "non-induced" samples. This permitted the later subtraction of the raw FIDs to remove the signal from the cellular background, which is due to the integration of the isotope-labeled amino acids in the proteome. The background broad signal from isotope-labeled peptides proved to be independent of the expression of the protein of interest, and thus can be removed using FID subtraction. However, the strong, sharper signals from abundant metabolites (e.g. amino acids) are often not cancelled perfectly. Also, because the number of cells in the tube was not always exactly the same, we had to apply a multiplication factor to the "non-induced" spectra to remove the cellular background signal. This factor varied between 0.8 and 1.2, which we fixed based on the best possible canceling of the baseline in the final spectrum. The subtraction of the "non-induced spectrum" was not necessary for $^{13}C$-Phe/$^{13}C$-Tyr, the cellular background signal being negligible. The addition and subtraction of raw FIDs were performed using Topspin 4. The peak intensity analysis was carried out using ccpNMR 3.1.1 [2].

*In vitro*, cell-extract and in-cell $^{1}H$-$^{15}N$ correlation spectra were obtained using two-dimensional $^{1}H$-$^{15}N$ HSQC (hsqcetfpf3gpsi2 from the Bruker library, $^{13}C$-decoupled in both dimensions) experiments at 10 °C, using an interscan delay of 0.6 s. We adjusted the number of acquired complex points and sweep widths to obtain similar FID resolutions. All $^{1}H$-$^{15}N$ HSQC spectra were processed using no-apodization and zero-filling to 4096 and 1024 points in the $^{1}H$ and $^{15}N$ dimensions, respectively.
For cell-extract and in-cell samples, $^{1}H$-$^{15}N$ HSQC NMR spectra were acquired with 2048 and 76 complex points, and sweep widths of 17 ppm and 14 ppm in the $^{1}H$ and $^{15}N$ dimensions, respectively. We recorded 1 hour-long spectra using 64 scans per $^{15}N$-increment. We verified that 4 consecutive spectra were superimposable, which revealed little evolution of our samples. Then we then added the raw FIDs of these 4 spectra to obtain a 4 hour-long spectrum.
*In vitro* reference $^{1}H$-$^{15}N$ HSQC NMR spectra of α-Syn were acquired with N-terminally acetylated [*u*-$^{13}C/^{15}N$]-α-Syn at 400 μM dissolved in PBS at pH 7.0. These NMR spectra were acquired with 1536 and 128 complex points and sweep widths of 16.23 ppm and 26 ppm in the $^{1}H$ and $^{15}N$ dimensions, respectively. We accumulated 4 scans per $^{15}N$-increment. DSS at ~100 μM and $D_2O$ at 3% (v/v) were added to these samples.
*In vitro* reference $^{1}H$-$^{15}N$ HSQC NMR spectra of tau were recorded with [*u*-$^{13}C/^{15}N$]-His6-tau at 55 μM in PBS at pH 7.0 at 283 K. These NMR spectra were acquired with 2048 and 792 complex points and sweep widths of 13.73 ppm and 24 ppm in the $^{1}H$ and $^{15}N$ dimensions, respectively. We accumulated 8 scans per $^{15}N$-increment. DSS at ~100 μM and $D_2O$ at 3% (v/v) were added to these samples.

All $^{13}C\alpha^{13}CO$ correlation spectra were acquired using the ($^{1}H$-flip*)-$^{13}Ca^{13}CO$-LB pulse sequence [3] at 37 °C. Pulses and delays were set up according to the parameters described in our previous publication. The pulse sequence is provided below. $^{13}C\alpha^{13}CO$ spectra were acquired with 1024 and 128 complex points, and sweep widths of 20.3 ppm and 14 ppm in the $^{13}CO$ and $^{13}C\alpha$ dimensions. The interscan delay was 0.2 s, which yields the best signal-to-noise ratio per unit of time for disordered proteins, according to our previous quantifications [3]. The total $^{13}C\alpha$ constant-time evolution was 27 ms. These $^{13}C\alpha^{13}CO$ spectra were acquired with 1024 and 128 complex points and sweep widths of 20.3 ppm and 14 ppm in the $^{13}CO$ and $^{13}C\alpha$ dimensions, respectively. This yielded acquisition times of 143 and 26 ms, and thus FID resolution of 7 and 38 Hz in $^{13}CO$ and $^{13}C\alpha$ dimensions, respectively. The $^{13}C\alpha^{13}CO$ spectra were processed with a square cosine bell and without apodization, and with zero-filling to 4096 and 1024 points in the $^{13}CO$ and $^{13}C\alpha$ dimensions, respectively. with 128 scans.
For in-extract and in-cell samples, we recorded 2 hour-long $^{13}C\alpha^{13}CO$ spectra using 128 scans per $^{13}C\alpha$-increment. We verified that 2 consecutive spectra were superimposable for α-Syn constructs, 4 consecutive spectra for tau, which revealed little evolution of our samples. Supplementary $^{1}H$-$^{13}C$ HSQC spectra were recorded before and after the $^{13}C\alpha^{13}CO$ acquisitions, which also revealed little evolution (a notable exception is to mention for the lactate signals). Then we then added the raw FIDs of these 2 or 4 spectra to obtain a 4 or 8 hour-long spectrum.
*In vitro* reference $^{13}C\alpha^{13}CO$ NMR spectra of α-Syn were acquired with N-terminally acetylated [*u*-$^{13}C/^{15}N$]-α-Syn at 200 μM dissolved in PBS at pH 7.0. We accumulated 4 scans per $^{13}C\alpha$-increment (~3.5 minutes-long acquisition). DSS at ~100 μM and $D_2O$ at 3% (v/v) were added to these samples.
*In vitro* reference $^{13}C\alpha^{13}CO$ NMR spectrum of tau was acquired with [*u*-$^{13}C/^{15}N$]-tau at 20 μM dissolved in PBS at pH 7.0. We accumulated 256 scans per $^{13}C\alpha$-increment (~4 hour-long acquisition). DSS at ~100 μM and $D_2O$ at 3% (v/v) were added to these samples.
The 950MHz-TCI probe had a S/N=1710 for $^{13}C$ ASTM (40% dioxane in $C_6D_6$, ASTM) when delivered. The 1200MHz-TCI-3mm probe had a S/N=805 for $^{13}C$ ASTM when delivered.

**Preparation of lipid vesicles**

Pig brain polar lipid extracts were purchased from Avanti. Lipid stock solutions were mixed in appropriate molar ratios in chloroform and lyophilized overnight. The dried lipids were solubilized in PBS (20 mM phosphate, 150 mM NaCl), pH 7.2. SUVs and LUVs were prepared by sonication (2 × 20 min, 30 % amplitude, Vibracell 75042 BIOBLOCK SC) on ice. LUVs were obtained in the case of pig brain polar lipid extract solubilized in PBS, while SUVs were obtained in many other buffers. We obtained SUVs from pig brain polar lipid extract solubilized and sonicated in hepes 20 mM, NaCl 50 mM, pH 7.2. Then, vesicles were centrifugated at 14,000 g to remove any metal residue from the sonicator probe. The vesicles size was determined by dynamic light scattering (DLS) (Nano series, Malvern).

##### Pulse sequence of ($^1$H-flip*)-$^{13}$Ca$^{13}$CO-LB

```
;c_hcaco_ctre_1_lb.ts4
;derived from c_hcaco_ctiare.2
;(H)CaCO
;2D sequence with
;   13C detected correlation for triple resonance using
;      multiple inept transfer steps
;
;      F2(Ha) -> F1(Ca,t1) -> F1(C=O,t2)
;
;on/off resonance 13C pulses using shaped pulses
;phase sensitive (t1)
;using constant time in t1
;relaxation optimised (H-flip)
;using 1 broadband 180 pulse (G5) in the final CA-CO transfer
;    use c_hcaco_ctre_2_lb.ts4 for 2 consecutive selective 180 pulses on CO and CA
;using low-bash scheme for 13CA real-time decoupling in t2
;
;A. Alik, C. Bouguechtouli, M. Julien, W. Bermel, R. Ghouil, S. Zinn-Justin & F.-X. Theillet,
;   Angew. Chem. 59, 10411-10415 (2020)
;W. Bermel, I. Bertini, I.C. Felli & R. Pierattelli,
;   J. Am. Chem. Soc. 131, 15340-5 (2009)
;(W. Bermel, I. Bertini, V. Csizmok, I. C. Felli, R. Pierattelli &
;   P. Tompa, J. Magn. Reson. 198, 275-281 (2009) )
;(L. Duma, S. Hediger, A. Lesage & L. Emsley,
;   J. Magn. Reson. 164, 187-195 (2003) )
;
;   for BASHD homodecoupling, read:
;J. Ying, F. Li, J.H. Lee, A. Bax,
;   J. Bioml. NMR 60, 15-21 (2014)
;
;   for G5 pulse, read:
;Y. Xia, P. Rossi, M.V. Subrahmanian, C. Huang, T. Saleh, C. Olivieri, C.G. Kalodimos, G. Veglia
;   J. Biomol. NMR 69, 237-243 (2017)
;   doi: 10.1007/s10858-017-0153-2
;
;$CLASS=HighRes
;$DIM=2D
;$TYPE=
;$SUBTYPE=
;$COMMENT=


prosol relations=<triple_c>

#include <Avance.incl>
#include <Grad.incl>
#include <Delay.incl>
#include <De.incl>

"p4=p3*2"
"d11=30m"
"d12=20u"
"d13=200u"

"d3=1.45m"
"d4=1.45m"
"d22=4.5m"
"d23=4.5m"
"d29=5m"

"d61=5m"
"d62=d61-p36-20u"
"d63=d62/2"
"l0=0.5*aq/d61+1"

"d0=3u"
"d20=d27-p12-4u"

"in0=inf1/2"
"in20=in0"

"DELTA2=p4+p25-p11"
"DELTA3=d3-p4/2"
"DELTA4=d3+p4/2"
"DELTA5=d22-3u-p3-p4"
"DELTA6=d27-d22-p12"

"cnst26=(cnst21+cnst22)/2"

"spoff21=0"
"spoff23=0"
"spoff24=0"
"spoff25=0"
```

```
"spoff26=bf1*((cnst21-cnst22)/1000000)"
"spoff28=0"
"spoff29=bf1*((cnst23-cnst22)/1000000)"
"spoff36=0"
"spoff52=bf1*((cnst26-cnst21)/1000000)"

1 ze
  d11 pl16:f3
  1m
  10u reset:f1
2 d11 do:f3
3 d1 fq=cnst22(bf ppm):f1
  d12 pl2:f2 cpd3:f3
  50u
  (p3 ph1):f2
  d4
  (center (p12:sp24 ph1) (p4 ph1):f2 )  ;pulse Q3 CA mid.select.
  d4
  (p3 ph2):f2
  DELTA2
  (p11:sp23 ph1)  ;pulse Q5 CA onres
  DELTA3
  (p4 ph2):f2
  (p23:sp21 ph1)  ;pulse Q3 CA high.select.
  DELTA4
  (p3 ph8):f2

  d0
  (p4 ph1):f2
  DELTA5
  (p12:sp26 ph7)  ;pulse Q3 CO offres
  DELTA6
  (p12:sp29 ph1)  ;pulse Q3 Cali mid.select.
  d20
  (p12:sp26 ph1)  ;pulse Q3 CO offres
  4u UNBLKGRAD
  (p11:sp25 ph4)  ;pulse Q5tr CA onres
  4u

;  p16:gp1
  d16 fq=cnst21(bf ppm):f1
  4u BLKGRAD

     (p11:sp25 ph5)  ;pulse Q5tr CO onres
     d23
     (p52:sp52 ph1)  ;pulse G5 CO-CA
     d23

 ACQ_START(ph30,ph31)
  0.1u START_NEXT_SCAN
  0.1u REC_UNBLK
  0.05u DWELL_RELEASE
  d63
  0.05u DWELL_HOLD
  0.1u REC_BLK

4 10u
  (p36:sp36 ph9):f1
  4u
  6u ipp9
  0.1u REC_UNBLK
  0.05u DWELL_RELEASE
  d62
  0.05u DWELL_HOLD
  0.1u REC_BLK
  10u
  (p36:sp36 ph9):f1
  4u
  6u ipp9
  0.1u REC_UNBLK
  0.05u DWELL_RELEASE
  d62
  0.05u DWELL_HOLD
  0.1u REC_BLK
  lo to 4 times l0

  0.1u REC_UNBLK
  0.05u DWELL_RELEASE
  d62
  0.05u DWELL_HOLD
  0.1u REC_BLK

  rcyc=2

  d11 do:f3 mc #0 to 2
  F1PH(calph(ph4, +90), caldel(d0, +in0) & caldel(d20, -in20))
  1m
```

```
exit

ph1=0
ph2=1
ph4=0 2
ph5=0 0 2 2
ph7=0 0 0 0 2 2 2 2
ph8=2
ph9=0 0 2 2 2 0 0 2 2 2 0 0 0 2 2 0
ph30=0
ph31=0 2 2 0

;pl1 : f1 channel - power level for pulse (default)
;pl2 : f2 channel - power level for pulse (default)
;pl16: f3 channel - power level for CPD/BB decoupling
;sp21: f1 channel - shaped pulse 180 degree  (CA on resonance, high selectivity, Q3_surbop)
;sp23: f1 channel - shaped pulse  90 degree  (on resonance, Q5_sebop)
;sp24: f1 channel - shaped pulse 180 degree  (CA on resonance, med. selectivity, Q3_surbop)
;sp25: f1 channel - shaped pulse  90 degree  (on resonance, Q5tr_sebop)
;                   for time reversed pulse
;sp26: f1 channel - shaped pulse 180 degree  (C=O off resonance, Q3_surbop)
;sp27: f1 channel - shaped pulse 180 degree  (CA off resonance, Q3_surbop)
;sp29: f1 channel - shaped pulse 180 degree  (Cali off resonance, Q3_surbop)
;sp36: f1 channel - inversion pulse for decoupling durinq acquisition (Sinc cosine-modulated at freq.
118ppm*bf1, C=O on resonance but side-band excitation at 54 and 290 ppm to compensate Bloch-Siegert shift)
;sp52: f1 channel - shaped pulse 180 degree  (G5 at <900 MHz, adiab.BIP720 at >1 GHz, C=O and Cali inversion,
off resonance centered at 113.75 ppm)
;p3 : f2 channel -  90 degree high power pulse
;p4 : f2 channel - 180 degree high power pulse
;p11: f1 channel -  90 degree shaped pulse         (231 usec at 700 MHz)
;p12: f1 channel - 180 degree shaped pulse         (med. selectivity, 300 usec at 700 MHz)
;p16: homospoil/gradient pulse                        [1 msec]
;p23: f1 channel - 180 degree shaped pulse         (high selectivity, 634 usec at 700 MHz)
;p36: f1 channel - 180 degree inversion during acquisition          (duration 22600/bf1, 129 usec at 700
MHz)
;p52: f1 channel - 180 degree shaped pulse (offset=113.75ppm, sp52, G5: length=5xp1 power=pl1, C=O and
Calibroad band inversion)
;d0 : incremented delay (F1 in 2D)                    [3 usec]
;d1 : relaxation delay                             [~200-400 msec]
;d3 : 1/(5J(HCa))                                     [1.45 msec]
;d4 : 1/(5J(HCa))                                     [1.45 msec]
;d11: delay for disk I/O                              [30 msec]
;d12: delay for power switching                       [20 usec]
;d16: delay for homospoil/gradient recovery        [200 usec]
;d20: decremented delay (F1 in 2D) = d27-p12-4u
;d22: 1/(4J(COCa))                                    [4.5 msec]
;d27: 1/(2J(CaCb) or 1/(J(CaCb)) or 3/(2J(CaCb))       [13.5 msec or 27 msec or 40.5msec]
;d61: length of total acquisition block            [5 msec]
;d62: length of block between decoupling pulses : = aq/l0
;d63: = d62/2
;cnst21: CO chemical shift (offset, in ppm)                [172 ppm]
;cnst22: Calpha chemical shift (offset, in ppm)            [55.5 ppm]
;cnst23: Caliphatic chemical shift (offset, in ppm) [41 ppm]
;o1p: CO chemical shift (cnst21)
;l0 : number of blocks during acquisition time
;       adjust to get d62 as required
;l6: flag fo mlev16 cycling during decoupling
;inf1: 1/SW(Ca) = 2 * DW(Ca)
;in0: 1/(2 * SW(Ca)) = DW(Ca)
;nd0: 2
;in20: = in0
;ns: 8 * n
;ds: >= 32
;td1: number of experiments in F1 ( < 4*d27*SWH1 )
;SW1(Ca) > 14 ppm
;SW2(CO) ~ 20 ppm
;FnMODE: States-TPPI (or TPPI) in F1
;cpd3: decoupling according to sequence defined by cpdprg3
;pcpd3: f3 channel - 90 degree pulse for decoupling sequence

;for z-only gradients:
;gpz1: 50%
;use gradient files:
;gpnam1: SMSQ10.100
```

**Table S1.** Amino acid concentration for isotope labeling.

| Amino acid | Concentration (mg/L) | Ref. supply |
|---|---|---|
| L-Asp.$H_2O$ | 10/15/70 | $^{13}C_4$, Eurisotop CLM-8699<br>$^{15}N_2$ Eurisotop NLM-3286 |
| L-Arg HCl | 84 | $^{13}C^{15}N$, Silantes 201604102 |
| L-Asn | 70 | $^{15}N_2$, CIL NLM-3286/ $^{13}C_4$, CIL CLM-8699 |
| L-Cys | 62 | $^{13}C_3{}^{15}N$, CIL CNLM-3871 |
| Gly | 120 | $^{15}N$ : Eurisotop N2451<br>1,2-$^{13}C$, Eurisotop CLM-1017 |
| L-His HCl | 42 | $^{13}C_6{}^{15}N_3$, Sigma-Aldrich 608009 |
| Ile precursor: Sodium-2-keto-3-methyl pentanoate | 105 | $^{13}C_6$, Eurisotop CLM-8669-PK |
| Leu precursor: $\alpha$-ketoisocaproic acid NaCl | 105 | 1,2 $^{13}C_2$, Eurisotop CLM-4826 |
| L-Lys.HCl | 146 | $^{13}C_6{}^{15}N_2$ Sigma-Aldrich, 608041 |
| L-Phe | 66 | $^{13}C^{15}N$, Silantes 213603900 |
| L-Tyr | 103 | $^{13}C^{15}N$, Silantes 218603900 |
| Val precursor $\alpha$-ketoisovaleric acid NaCl | 94 | $^{13}C_5$,Eurisotop CLM-4418 |

**Table S2:** $^{1}$H-$^{13}$C crosspeak intensities in HSQC spectra of in-cell samples supplemented with single $^{13}$C-labeled amino acids (Leu is only 1,2-$^{13}$C-labeled). Intensities are normalized according to $^{1}$H-$^{13}$C HSQC spectra from natural abundance, non-induced cells, which contain high concentrations of natural abundance amino acids. The surrounding DMEM+FBS in the tube contains also natural abundance amino acids at constant concentrations in the induced and non-induced samples.

| isotope labeling / assigned peaks | | Arg | Asn | Asp | Cys | Gly | Ile | Leu | Lys | Phe/Tyr | Val |
|---|---|---|---|---|---|---|---|---|---|---|---|
| Ala | CαHα | 0.5 | 1.1 | 1.3 | 0.3 | 1.6 | 1.3 | 0.5 | 0.6 | 1.4 | 0.4 |
| | CβHβ | 0.5 | 1.1 | 1.6 | 0.4 | 1.4 | 1.4 | 0.6 | 0.6 | 1.5 | 0.5 |
| Arg | Cβ | 17.8 | 1.7 | 1.1 | 1.2 | 1.3 | 1.0 | 0.8 | 1.6 | 1.9 | 1.0 |
| | Cγ | 8.6 | 0.4 | 0.3 | 0.6 | 1.1 | 1.0 | 0.9 | 1.8 | 1.0 | 0.6 |
| | Cγ | 18.2 | 0.8 | 0.6 | 0.9 | 0.9 | 1.0 | 2.9 | 1.4 | 1.1 | 0.6 |
| | C.Hδ | 31.8 | 0.7 | 0.4 | 1.1 | 0.9 | 0.6 | 0.8 | 1.2 | 0.8 | 0.7 |
| Unassigned | ? | 69.6 | 3.2 | 1.8 | 1.8 | 3.0 | 1.9 | 0.6 | 5.6 | 4.5 | 0.3 |
| Asn | CαHα | -0.1 | 40.2 | 0.6 | 0.1 | -1.2 | 0.3 | 0.2 | 0.0 | 0.3 | 0.2 |
| | CβHβ | 0.2 | 47.5 | 5.2 | 0.1 | 0.5 | 0.8 | 0.7 | 1.7 | 0.1 | 0.3 |
| Asp | CβHβ | 0.8 | 21.2 | 128.1 | 0.9 | 0.8 | 0.6 | 1.0 | 0.8 | 1.4 | 0.6 |
| Glutathione | red. (Cys moiety) | 0.3 | 0.5 | 0.5 | 9.6 | 0.8 | 0.5 | 0.6 | 2.5 | 0.9 | 0.5 |
| | ox. (Cys moiety) | 1.2 | 0.9 | 0.5 | 1.1 | 1.0 | 0.8 | 1.0 | 36.3 | 1.0 | 0.9 |
| | red+ox (Gly moeity) | 0.3 | 0.6 | 0.9 | 0.4 | 47.5 | 0.6 | 0.5 | 0.3 | 0.9 | 0.5 |
| Gly | CαHα | 1.1 | 1.4 | 1.8 | 2.6 | 180.2 | 2.1 | 2.4 | 1.1 | 1.9 | 3.0 |
| Unassigned | ? | 0.8 | 0.6 | 0.7 | 1.3 | 48.3 | 0.8 | 1.2 | 1.1 | 1.1 | 1.1 |
| Unassigned | ? | 0.7 | 1.2 | 0.9 | 1.0 | 68.4 | 0.6 | 1.7 | 1.5 | 1.0 | 1.0 |
| Gln | Cβ-Hβ | 1.0 | 0.6 | 1.1 | 0.3 | 1.4 | 0.3 | 0.4 | 0.6 | 0.5 | 0.2 |
| Glu | Cβ-Hβ | 0.5 | 0.7 | 0.9 | 0.4 | 0.9 | 0.8 | 0.6 | 0.6 | 0.8 | 0.5 |
| Pro | CαHα | 5.1 | 3.2 | 3.6 | 1.5 | 1.3 | 2.9 | 1.5 | 1.7 | 2.3 | 1.5 |
| | Cδ-Hδ1 | 5.3 | 1.9 | 1.9 | 1.1 | 0.8 | 2.3 | 2.2 | 1.3 | 1.8 | 1.0 |
| | Cδ-Hδ2 | 4.6 | 1.5 | 1.6 | 0.9 | 0.7 | 2.0 | 2.0 | 0.8 | 1.7 | 1.2 |
| | Cγ Hγ | 6.5 | 1.0 | 0.7 | 1.3 | 0.8 | 0.8 | 0.9 | 0.9 | 0.9 | 0.8 |
| | CβHβ1 | 4.4 | 2.8 | 3.7 | 1.2 | 1.1 | 2.1 | 1.3 | 1.0 | 1.6 | 1.9 |
| | CβHβ2 | 4.7 | 1.9 | 3.1 | 1.0 | 0.9 | 1.9 | 0.6 | 0.7 | 1.4 | 1.1 |
| Ile | $CH_3$ | 1.3 | 8.0 | 1.1 | 1.0 | 1.5 | 15.9 | 0.6 | 0.3 | 1.5 | 0.9 |
| | C.H | 1.3 | 5.6 | 0.5 | 1.0 | 1.5 | 10.9 | 0.8 | 1.1 | 1.3 | 1.0 |
| Leu | Cδ Hδ1 | 1.0 | 0.9 | 0.8 | 1.2 | 1.2 | 1.1 | 1.4 | 1.4 | 1.1 | 1.0 |
| | Cδ. Hδ2 | 1.3 | 0.9 | 1.0 | 1.0 | 1.3 | 1.4 | 1.1 | 1.6 | 1.3 | 1.8 |
| | CβHβ1 | 0.5 | 0.7 | 0.6 | 0.9 | 0.8 | 0.6 | 0.6 | 6.4 | 0.8 | 0.7 |
| | CβHβ2 | 0.6 | 1.2 | -0.1 | 1.2 | 1.3 | 1.0 | -0.2 | 7.4 | 1.0 | 0.2 |
| Lys | CεHε | 1.2 | 0.9 | 0.5 | 1.1 | 1.0 | 0.8 | 1.0 | 36.3 | 1.0 | 0.9 |
| | CγHγ1 | 0.7 | 0.8 | 0.5 | 1.0 | 0.8 | 0.7 | 1.1 | 31.6 | 0.8 | 1.1 |
| | CγHγ2 | 1.1 | 0.9 | 0.6 | 1.4 | 1.0 | 0.9 | 1.6 | 41.0 | 1.2 | 1.0 |
| | CδHδ | 0.9 | 0.8 | 0.6 | 1.2 | 1.0 | 0.8 | 1.0 | 29.6 | 1.0 | 1.0 |
| | CβHβ | 1.5 | 0.9 | 0.6 | 0.9 | 1.1 | 0.8 | 1.1 | 19.0 | 1.0 | 2.0 |
| unassigned | | 1.0 | 0.6 | 0.5 | 1.2 | 1.1 | 0.8 | 1.3 | 37.4 | 1.2 | 1.7 |
| | | 1.2 | 0.6 | 0.6 | 1.3 | 1.1 | 0.9 | 1.3 | 36.3 | 1.0 | 2.0 |
| Phe | CβHβ1 | 3.7 | 1.6 | -0.1 | 0.9 | 1.5 | 2.2 | 1.4 | 1.4 | 9.5 | 0.6 |
| | CαHα | 0.6 | 0.3 | 0.1 | 1.1 | 1.1 | 1.1 | 0.2 | 4.2 | 4.5 | 1.3 |
| Ser | CβHβ | 0.9 | 1.2 | 1.4 | 1.4 | 0.3 | 1.2 | 1.7 | 0.7 | 1.6 | 1.7 |
| Tyr | CβHβ1 | 1.4 | -0.2 | 0.2 | 0.2 | 0.8 | -0.2 | 0.5 | -0.7 | 5.7 | 0.1 |
| Unassigned | val prec ? | 1.1 | 0.8 | 0.6 | 1.2 | 1.2 | 0.8 | 1.2 | 1.2 | 1.6 | 44.9 |
| Val | CαHα | 0.7 | 1.7 | 2.1 | 1.4 | 1.6 | 2.5 | 3.1 | 0.9 | 2.5 | 4.8 |
| | CβHβ | 5.4 | 1.2 | 1.9 | 1.1 | 1.1 | 1.1 | 0.5 | 1.0 | 1.2 | 4.9 |
| | Cγ1Hγ | 1.3 | 1.1 | 0.9 | 0.9 | 1.5 | 1.5 | 1.2 | 1.3 | 1.7 | 16.9 |
| | Cγ2Hγ | 1.2 | 1.1 | 1.4 | 1.4 | 1.7 | 2.1 | 1.6 | 1.5 | 2.1 | 19.1 |
| ATP | C1H1 | 0.9 | 1.1 | 0.7 | 1.4 | 1.2 | 0.7 | 1.7 | 0.7 | 1.0 | 1.9 |
| | C1H2 | 1.3 | 1.0 | 0.5 | 1.6 | 1.2 | 0.4 | 1.4 | 0.6 | 1.0 | 1.3 |
| Lactate | C1 | 1.0 | 1.1 | 0.8 | 1.1 | 1.0 | 1.0 | 1.0 | 1.1 | 1.0 | 1.0 |
| | C2 | 1.0 | 1.0 | 1.0 | 1.0 | 0.9 | 1.0 | 1.0 | 1.0 | 1.0 | 1.0 |

**Table S3.** Amino acid specific labelling for in-cell NMR.

| Amino acid | $^{15}N$ | $^{13}C$ |
|---|---|---|
| Asp | NO | NO |
| Arg | YES ? | YES + Proline |
| Asn | YES | NO |
| Cys | NO | NO |
| Gly | YES – Ser + glutathione | YES – Ser + glutathione |
| His HCl | YES ? | YES |
| Ile | NO | YES |
| Leu | NO | YES |
| Lys.HCl | YES | YES |
| Phe | YES | YES |
| Tyr | YES | YES |
| Val | NO | YES |

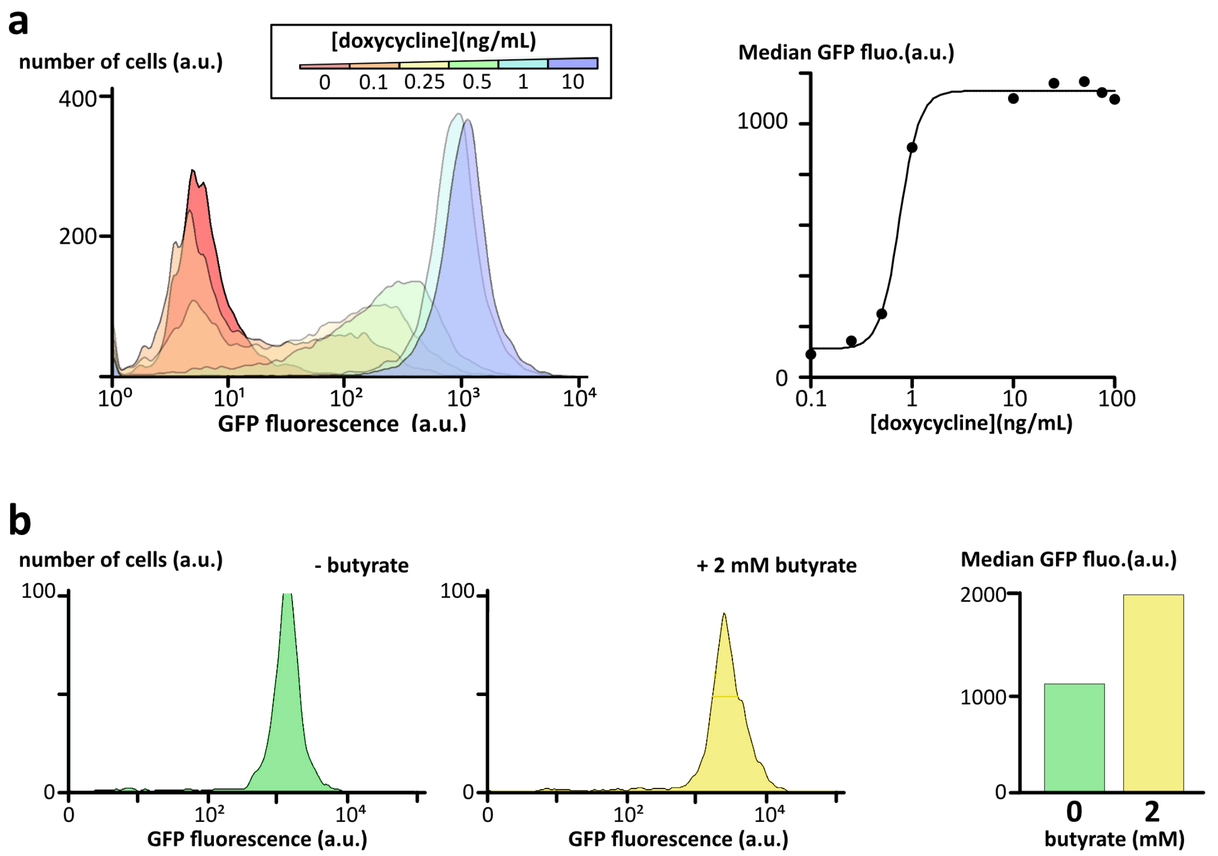


***Figure S1: a)** Expression levels of the chimera EGFP-$\alpha$-syn using fluorescence cell cytometry in stable inducible HEK Flp-In T-Rex cell lines in function of doxycycline concentration; **b)** Expression levels of EGFP-$\alpha$-syn at 10 ng/mL of doxycycline in presence (yellow) or absence (green) of sodium butyrate at 2 mM.*

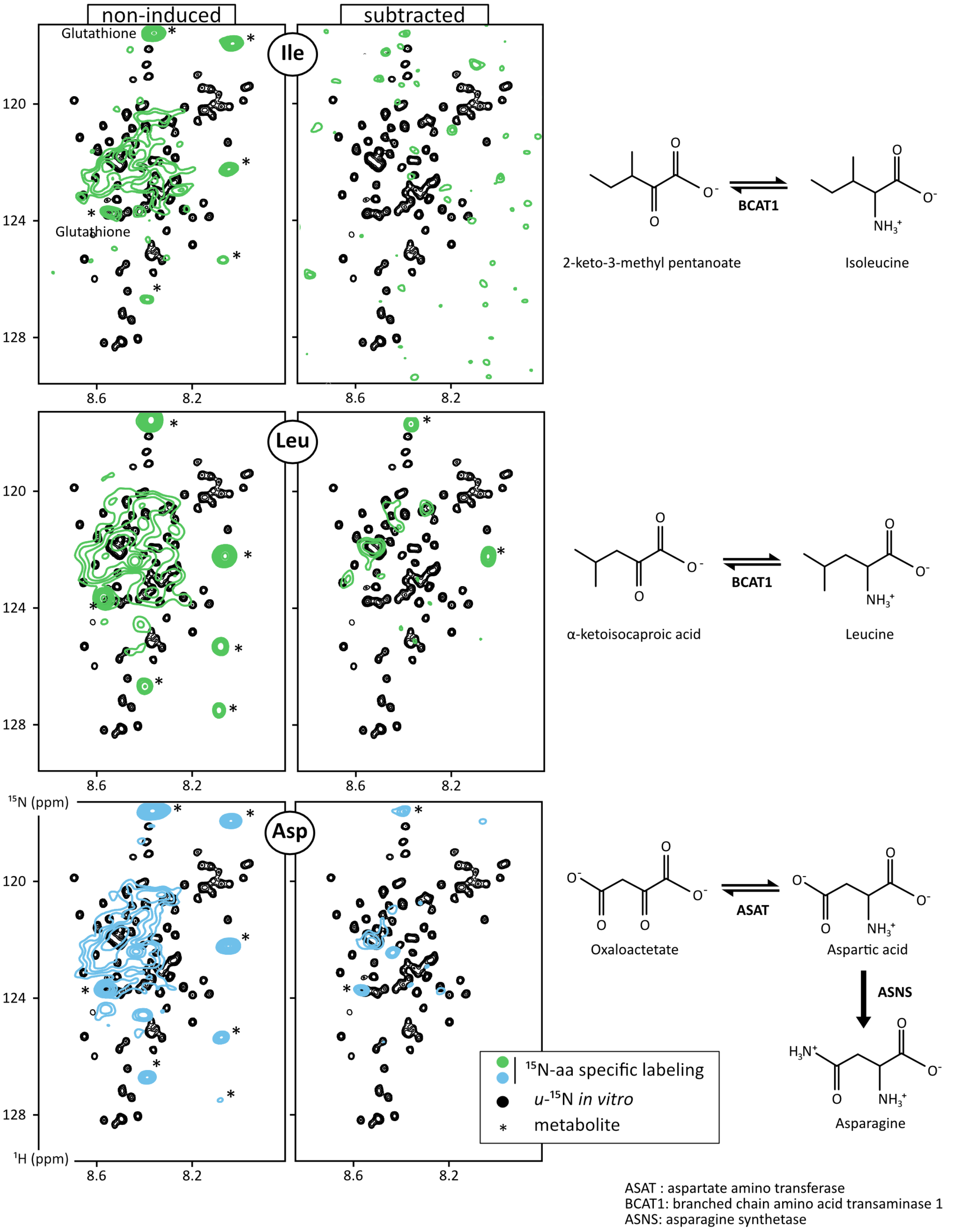


**Figure S2:** Overlays of $^{1}$H-$^{15}$N HSQC spectra of purified α-Syn (black) and in-cell spectra recorded from non-induced cells (left) or from cells expressing α-Syn (right, after subtraction of non-induced spectra) in a culture medium supplemented with $^{15}$N-Leu, $^{15}$N-Val or $^{15}$N-Asp. The semi-developed chemical structures are those of the amino acids and their immediate keto-acids. The known active transaminases are indicated.

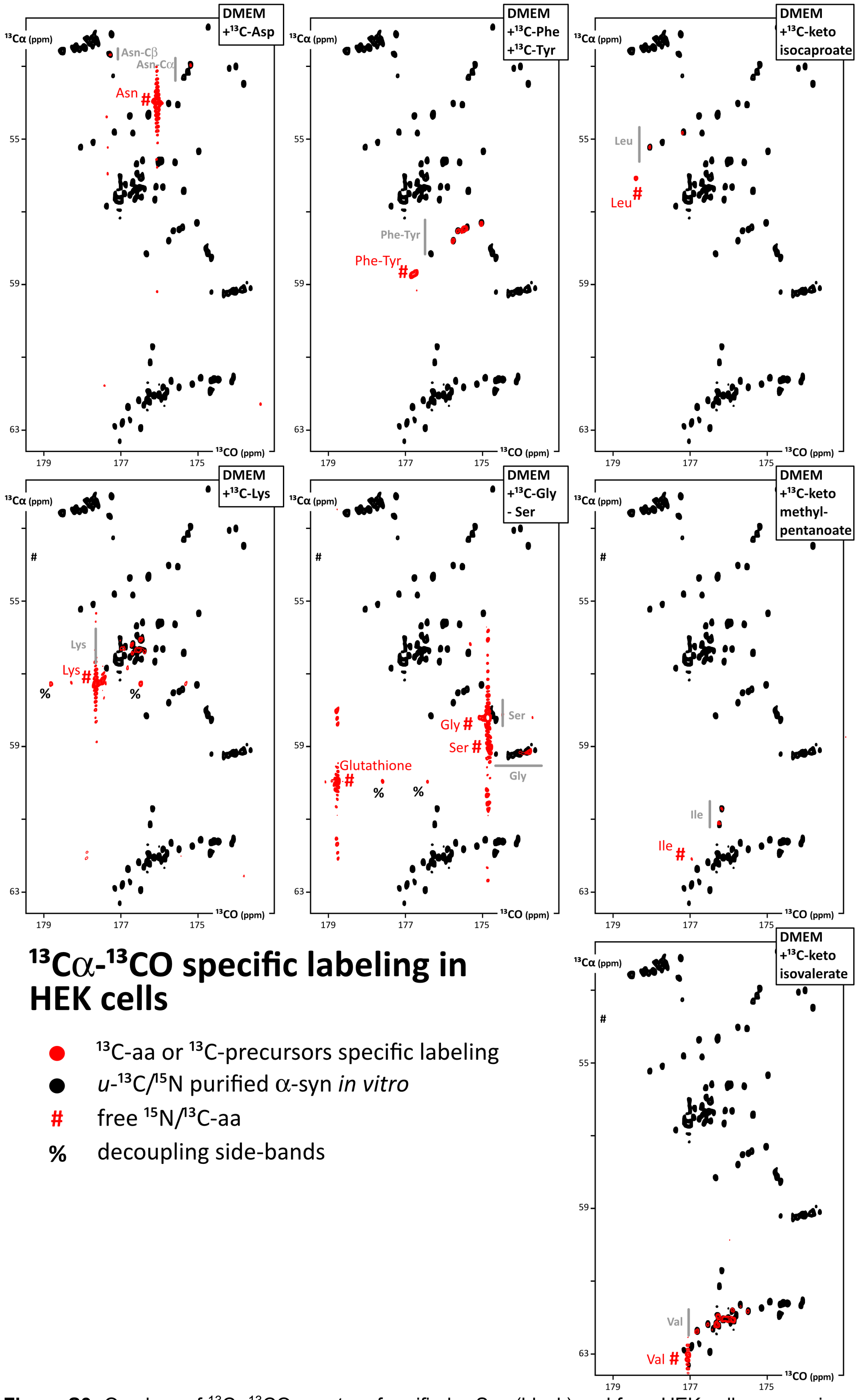


**Figure S3:** Overlays of $^{13}C\alpha^{13}CO$ spectra of purified α-Syn (black) and from HEK cells expressing α-Syn (red) in presence of $^{13}C$-Asp, or -Lys, or -Phe/Tyr, or -Gly, or $^{13}C$-ketoisocaproate, or $^{13}C$-ketomethylbutanoate, or $^{13}C$-ketoisovalerate. These amino acids or precursors thereof were supplemented in a home-made DMEM medium instead of their non-labeled counterparts, using the quantities mentioned in Table S1.

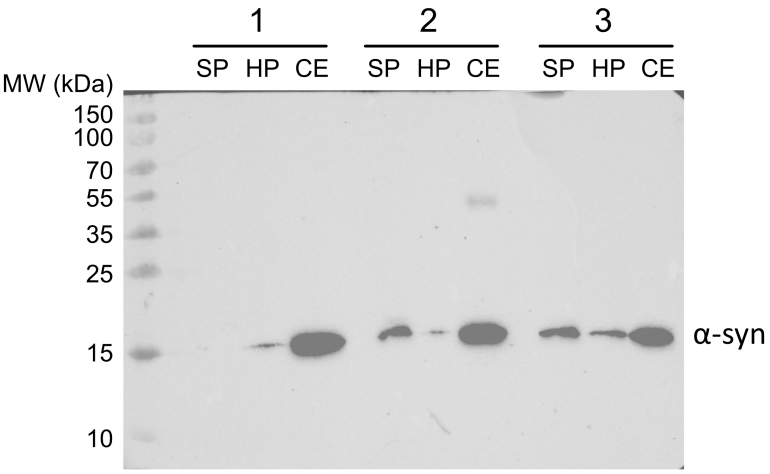


**Figure S4:** Western-blot analysis (3 independent samples) of a-syn in lysates from HEK-α-Syn (Flp-In TRex cells): pellets after sonication (SP), pellets after heating 3 min at 95 °C (HP), and soluble fraction after heating 3 min at 95 °C (clarified extract, CE).

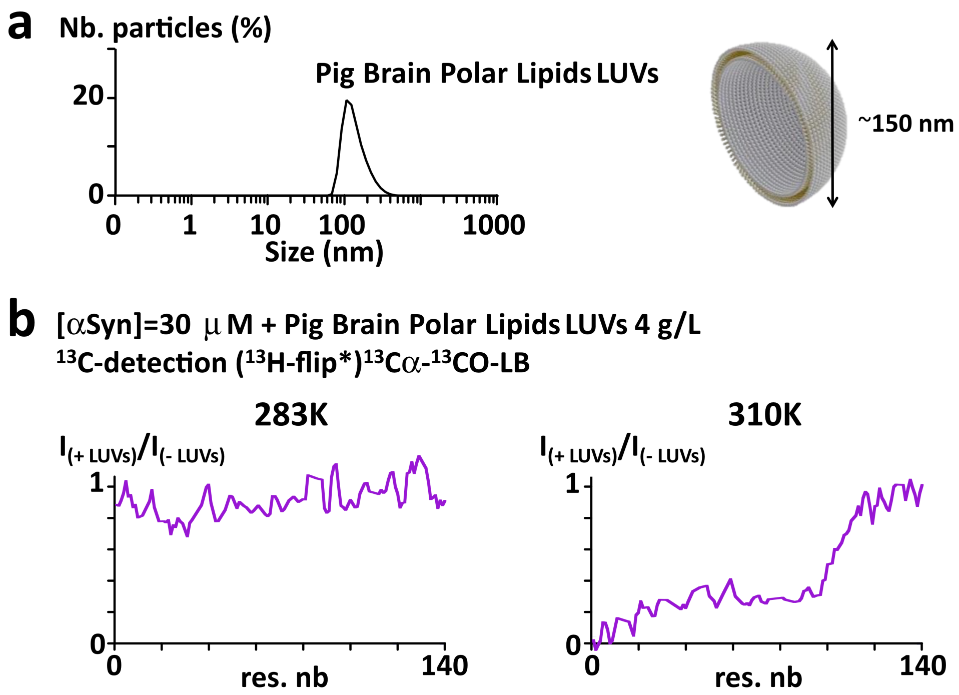


**Figure S5: a)** DLS profile of the vesicle population obtained from Pig Brain Polar Lipids sonicated in PBS. **b)** Residue specific intensity ratios of α-Syn in $^{13}$Cα $^{13}$CO spectra recorded in presence vs absence of large unilamellar vesicles (LUVs) made of Pig Brain Polar Lipids at 283 K (left) or 310 K (right). Peak disappearance reveals membrane binding only at 310 K.

These data were obtained using a $^{13}$C-low-sensitivity probe ($^{1}$H[$^{13}$C/$^{15}$N] TCI Bruker from 2006: $^{13}$C ASTM S/N=850) on a 700 MHz Bruker Avance Neo spectrometer.

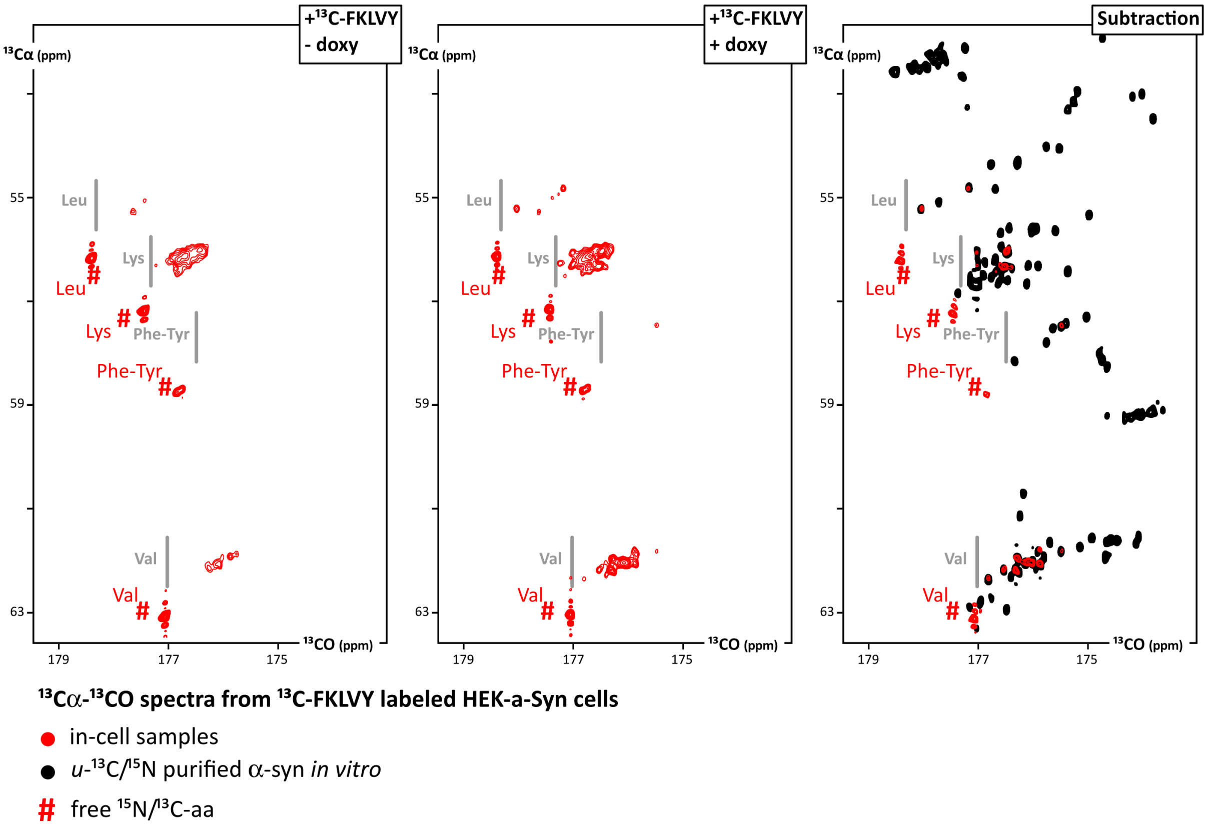


**Figure S6:** $^{13}C\alpha^{13}CO$ spectra of HEK(FlpIn TRex)-α-Syn cells cultured 48h in DMEM containing $^{13}C$-FKY and $^{13}C$-ketoisocaproate/ketoisovalerate in absence of doxycycline (left), in presence of doxycycline (middle) and the subtraction of these spectra (right, in red) overlayed with the spectrum of purified *u*-$^{13}C$-α-Syn (black).

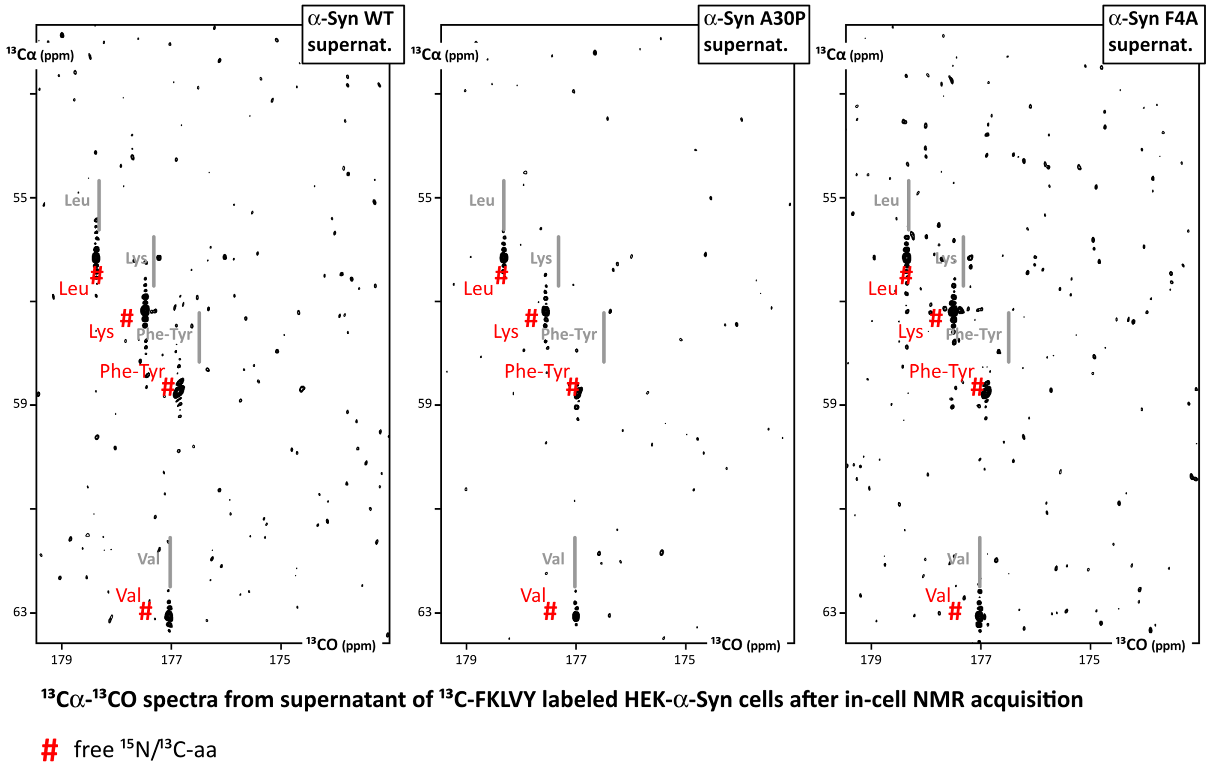


**Figure S7:** $^{13}$C$\alpha^{13}$CO spectra of the supernatant recovered from in-cell samples after acquisition, resuspended and pelleted by centrifugation. The 3 spectra were acquired in 4 hours using the same tubes and NMR parameters; they can thus be directly compared with the in-cell NMR spectra (see Figure S6). The contour level was set at the emergence of the background noise and shows no visible peaks from peptides; the peaks from the labeled free amino acids show that the plasmid membranes are permeable for amino acids.

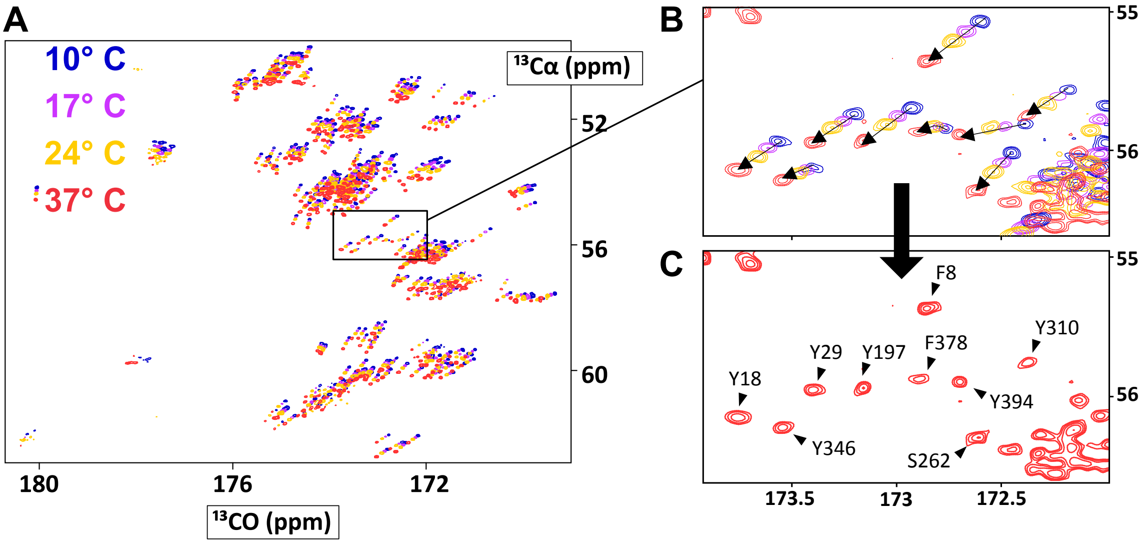


**Figure S8:** Transfer of peak assignment for tau-2N4R from 10 to 37 °C. **A.** Overlay of $^{13}C\alpha$-$^{13}CO$ spectra recorded at 10, 17, 24 et 37 °C with $^{15}N$-$^{13}C$ tau at 55 μM in PBS pH 7.0 (700 MHz, TXO). **B.** Close-up view in the region of Tyr/Phe. **C.** Assignment of the $^{13}C\alpha$-$^{13}CO$ spectrum of tau at 37 °C.

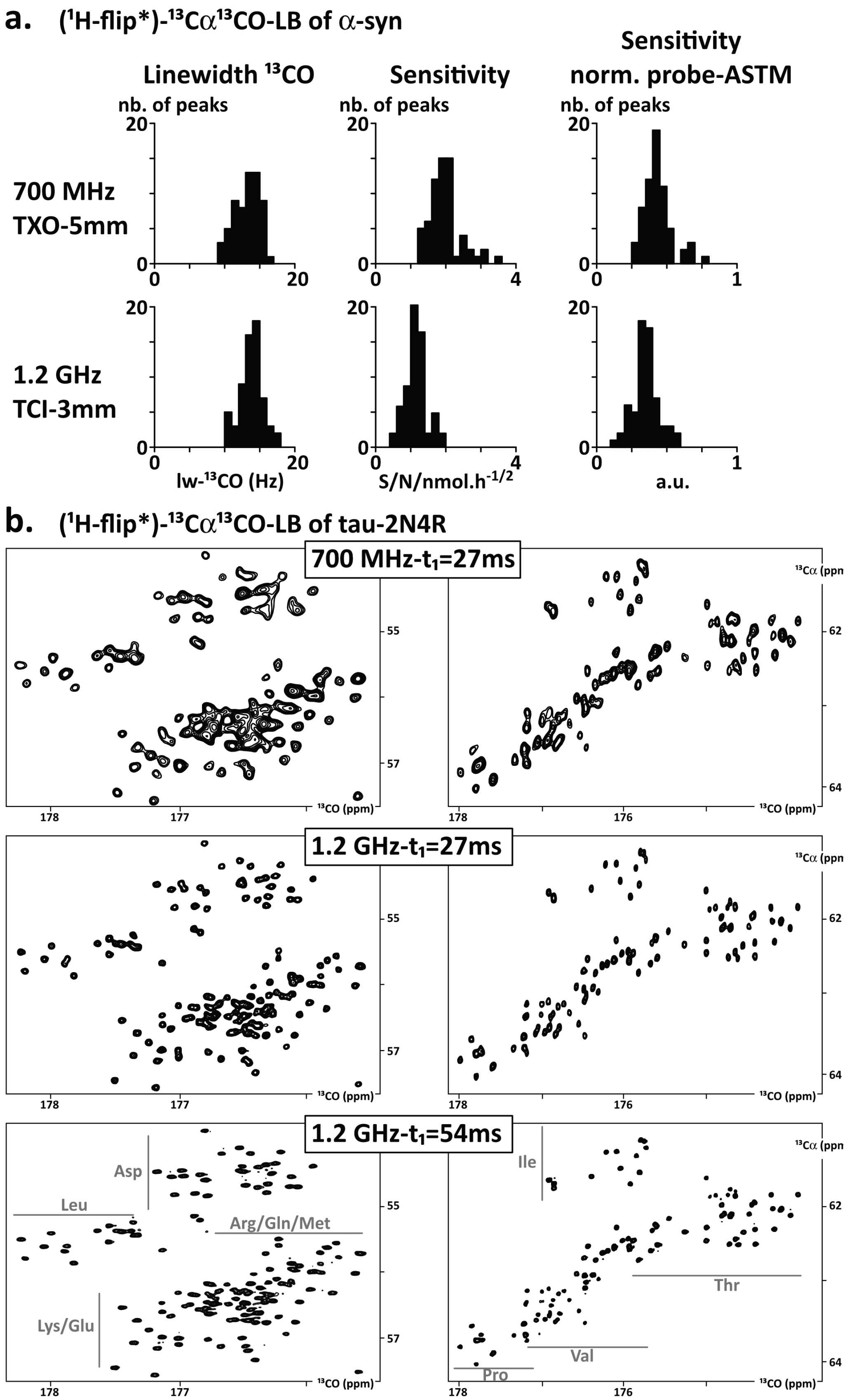


**Figure S9: a)** Distributions among 60 isolated peaks of α-syn (*in vitro*) of $^{13}$CO linewidths, sensitivity, and sensitivity divided by the probe $^{13}$C-ASTM sensitivity at 700 MHz and 1.2 GHz. Spectra were acquired with 3 mm tubes. An excellent signal linearity is observed from 3 to 5 mm in PBS. **b)** Close-up views of $^{13}$Cα-$^{13}$CO spectra of purified tau recorded at 700 MHz (black) or at 1.2 GHz (red and blue). *Left:* region of Leu/Lys/Asp/Arg/Gln/Glu; *Right:* region of Pro/Val/Thr/Ile. The constant-time evolution in the indirect dimension was set to 27 or 54 ms.
